# SMORE: joint dimension reduction and cell population discovery on single-cell methylome data

**DOI:** 10.64898/2026.09.14.751514

**Authors:** Jingwen Deng, Zixi Wang, Wen Tang, Guanyu Hu, Hao Feng

## Abstract

Single-cell DNA methylation profiling technology captures novel epigenetic data modality but are challenging to analyze because of their heterogeneity, high dimensionality, and ultra-sparsity. Here we present SMORE (*S*ingle-cell *M*ethyl*O*me *R*eduction and *E*mbedding), a computational method for joint dimensionality reduction and cell population discovery dedicated to single cell DNA methylation data. SMORE operates on a Bayesian framework that converts methylation proportions into ordered methylation states and jointly infers a low-dimensional representation, cell populations and their number. By using low-rank latent Gaussian factorization and adopting a mixture-of-finite-mixtures prior on latent cell scores, SMORE infers cell assignments without requiring a prespecified cluster number, and propagates uncertainty from methylation measurements to cell assignments. Across simulations spanning varying sample sizes, population imbalance, signal strengths and model misspecification, *SMORE* accurately recovered latent population structure and outperformed existing methods. Applied to human single-cell methylation datasets from lung, peripheral blood and primary motor cortex, *SMORE* recovered biologically supported cell population structures. *SMORE* provides an uncertainty-aware framework for dimension reduction and population discovery for single-cell methylomes.

## 1 Introduction

Rapid advances in single-cell and single-nucleus DNA methylation (scDNAm) profiling have transformed the study of epigenetic regulation from population-averaged measurements to cell-resolved maps of the methylome. Building on early single-cell bisulfite-sequencing protocols (Smallwood et al., 2014; Farlik et al., 2015), improvements in molecular barcoding, combinatorial indexing, sequencing throughput and joint multi-omic profiling now enable DNA methylation to be measured across thousands to tens of thousands of individual cells, often alongside transcriptomic, chromatin-accessibility or three-dimensional genome information (Angermueller et al., 2016; Luo et al., 2017; Mulqueen et al., 2018; Clark et al., 2018; Luo et al., 2018; D.-S. Lee et al., 2019). In particular, the development of single-nucleus methylcytosine sequencing and its subsequent refinements, including snmC-seq2 and snmC-seq3, substantially improved library complexity, genome coverage and experimental throughput, facilitating the construction of increasingly comprehensive cell-type-resolved methylome atlases (Luo et al., 2017, 2018; Liu et al., 2021, 2023). Combinatorial-indexing and droplet-based strategies further increased the number of cells that could be processed in parallel, whereas multi-omic protocols enabled DNA methylation to be jointly examined with gene expression, chromatin accessibility and three-dimensional chromatin organization in the same cells (Angermueller et al., 2016; Mulqueen et al., 2018; Clark et al., 2018; D.-S. Lee et al., 2019). These technologies have revealed extensive epigenetic heterogeneity obscured in bulk samples and have provided new insights into rare cell populations and lineage-associated cell states (Farlik et al., 2015; Luo et al., 2017; Liu et al., 2021, 2023), embryonic development (Zhu et al., 2018), aging (Hernando-Herraez et al., 2019), and tumor heterogeneity and evolution (Johnson et al., 2021). More recently, these efforts have culminated in the first body-wide human single-cell atlas jointly profiling three-dimensional genome organization and DNA methylation (Zhou et al., 2026), providing an important community resource for investigating the epigenomic basis of gene regulation, cell differentiation and human disease.

At the same time, the extreme sparsity, uneven genomic coverage and binary nature of single-cell methylation measurements create analytical challenges that differ fundamentally from those encountered in single-cell transcriptomics. A growing computational ecosystem has therefore emerged for methylation-state imputation, feature aggregation, dimensionality reduction, clustering, cell-type annotation, trajectory reconstruction, multi-omic integration and differential methylation analysis (Iqbal & Zhou, 2023). For methylation-state imputation, methods such as DeepCpG use deep neural networks to borrow information from local CpG neighborhoods and genomic sequence, whereas Melissa and Epiclomal employ probabilistic models that jointly infer missing methylation states and cellular population structure (Angermueller et al., 2017; Kapourani & Sanguinetti, 2019; de Souza et al., 2020). Feature aggregation over genomic regions, genes and regulatory elements is commonly used to reduce sparsity before dimensionality reduction. Specialized frameworks address complementary components of this workflow: scMET identifies highly variable features and supports differential methylation and variability analyses (Kapourani et al., 2021), whereas epiScanpy and ALLCools provide functionality for feature processing, low-dimensional representation and graph-or cluster-based analysis of single-cell methylation data (Danese et al., 2021; Liu et al., 2021). Trajectory reconstruction and multi-omic integration have likewise become important components of scDNAm analysis. MATCHER aligns pseudotemporal trajectories inferred from separately measured transcriptomic and epigenetic data, whereas LIGER and related latent-factor approaches, including MOFA+, integrate methylation with transcriptomic or chromatin-accessibility measurements in shared low-dimensional representations (Welch et al., 2017, 2019; Argelaguet et al., 2020; Iqbal & Zhou, 2023). For differential methylation analysis, scMET models feature-specific methylation levels and cell-to-cell variability, enabling the identification of differential mean methylation and differential variability across biological conditions (Kapourani et al., 2021). More recently, the mist was developed to model DNA methylation trajectories along a supplied pseudotime and to identify genomic features exhibiting temporal changes or phenotype-specific differences in their methylation dynamics (Duan et al., 2026).

Dimensionality reduction and clustering are foundational to single-cell analysis because they transform high-dimensional molecular profiles into representations that reveal cell–cell relationships, support visualization and annotation, and enable the discovery of rare, transitional or previously unrecognized cell populations. Early analyses of scDNAm data typically aggregated sparsely observed CpGs into genomic windows or functional regions, and then applied general-purpose techniques such as principal component analysis (PCA) (Greenacre et al., 2022), non-negative matrix factorization (NMF) (D. D. Lee & Seung, 1999), or uniform manifold approximation and projection (UMAP) (McInnes et al., 2018), followed by clustering. For example, epiScanpy implements feature construction, dimensionality reduction, nearest-neighbor graph analysis and community detection for single-cell epigenomic data (Danese et al., 2021). Such workflows are widely accessible and can perform well when populations are clearly separated, but the inferred partition may depend on feature selection, imputation, distance metrics, embedding parameters and the choice of clustering resolution or cluster number (Liu et al., 2023). Moreover, methods developed for continuous measurements effectively treat numerically coded methylation states as equally spaced, an assumption that need not hold for ordered categories derived from noisy or sparsely covered proportions (Korhonen et al., 1998; Choi, 2010). Methylome-dedicated approaches subsequently emerged, including pairwise-dissimilarity methods such as PDclust (Hui et al., 2018); Bayesian mixture models such as Melissa and Epiclomal, which borrow information across cells and genomic loci while clustering cells (Kapourani & Sanguinetti, 2019; de Souza et al., 2020); scMelody, which combines multiple cell–cell similarity measures through consensus clustering (Tian et al., 2022); and MethSCAn, which identifies informative genomic regions for exploratory representation (Kremer et al., 2024). More recently, MethylVI and scMethCraft have used latent-variable or neural-network models to learn cell embeddings while accounting for technical variation or incorporating genomic sequence and intercellular similarity (Weinberger et al., 2026; Tang et al., 2026).

Despite this progress, important methodological gaps remain. General-purpose workflows do not explicitly accommodate the binary-at-locus, bounded-after-aggregation and coverage-dependent nature of scDNAm measurements, whereas extensive binning, filtering or imputation can obscure focal methylation signals and rare cellular states (Iqbal & Zhou, 2023; Kremer et al., 2024; Weinberger et al., 2026). Existing specialized methods also entail trade-offs among modeling resolution, scalability and interpretability and can be sensitive to feature selection, similarity definitions, initialization or prespecification of the cluster structure (Kapourani & Sanguinetti, 2019; de Souza et al., 2020; Tian et al., 2022; Iqbal & Zhou, 2023). Moreover, dimensionality reduction and clustering are commonly treated as separate objectives, such that the resulting embedding is not explicitly optimized to recover stable and biologically meaningful cell populations (Danese et al., 2021; Kremer et al., 2024; Weinberger et al., 2026). A scalable statistical framework that jointly learns methylation-aware low-dimensional representations and cellular clusters while propagating uncertainty from sparse observations therefore remains needed. Here, we introduce *SMORE* (Single-cell MethylOme Reduction and Embedding), a Bayesian computational framework for jointly learning low-dimensional representations and identifying cell populations from single-cell methylome data. *SMORE* models observed methylation levels as ordinal states and links them to underlying continuous latent variables through a data-augmentation scheme (Albert & Chib, 1993). A low-rank factor model captures coordinated methylation variation across genomic regions and yields cell-specific coordinates that define a shared latent embedding. A mixture-of-finite-mixtures prior on the latent factor scores enables *SMORE* to infer both cell assignments and the number of occupied populations, eliminating the need to prespecify the number of clusters (Miller & Harrison, 2018). Posterior inference is summarized using pairwise co-clustering probabilities and a representative partition, allowing uncertainty in cell membership and population structure to be quantified directly. By unifying the ordinal observation model, latent representation and cluster discovery within a single probabilistic formulation, *SMORE* propagates uncertainty throughout the analysis instead of treating dimensionality reduction and clustering as separate analytical stages.

We benchmarked *SMORE* using two complementary simulation frameworks and human single-cell DNA methylation datasets from lung, peripheral blood and primary motor cortex. Model-based simulations evaluated recovery under the assumed ordinal latent-factor mixture structure, whereas empirically calibrated simulations reproduced data characteristics estimated from an independent single-nucleus methylation dataset. Across both model-based simulation and real-data-based simulation settings, *SMORE* accurately recovered the underlying population structure and remained robust to variation in signal strength and population imbalance. In real datasets, *SMORE* generated informative low-dimensional representations and cell partitions that closely agreed with reference cell-type annotations across diverse tissues, although the magnitude of its relative gains varied with the composition of the data set. These results demonstrate the ability of *SMORE* to resolve cellular heterogeneity from sparse single-cell methylation profiles while quantifying uncertainty in population number and cell assignments.

## 2 Results

### 2.1 Overview of *SMORE*

*SMORE* analyzes a preprocessed methylation matrix in which rows correspond to genomic regions and columns to individual cells (Fig. 1A). The methylation levels are converted from continuous observations between 0 and 1 to an interpretable ordinal methylation state. Its output is a posterior distribution over cell partitions rather than a single clustering obtained after a fixed embedding. The framework connects the observed methylation states, a low-dimensional representation of the cells and clustering discovery within one Bayesian model, allowing uncertainty to propagate across the complete analysis.

**Figure 1:**
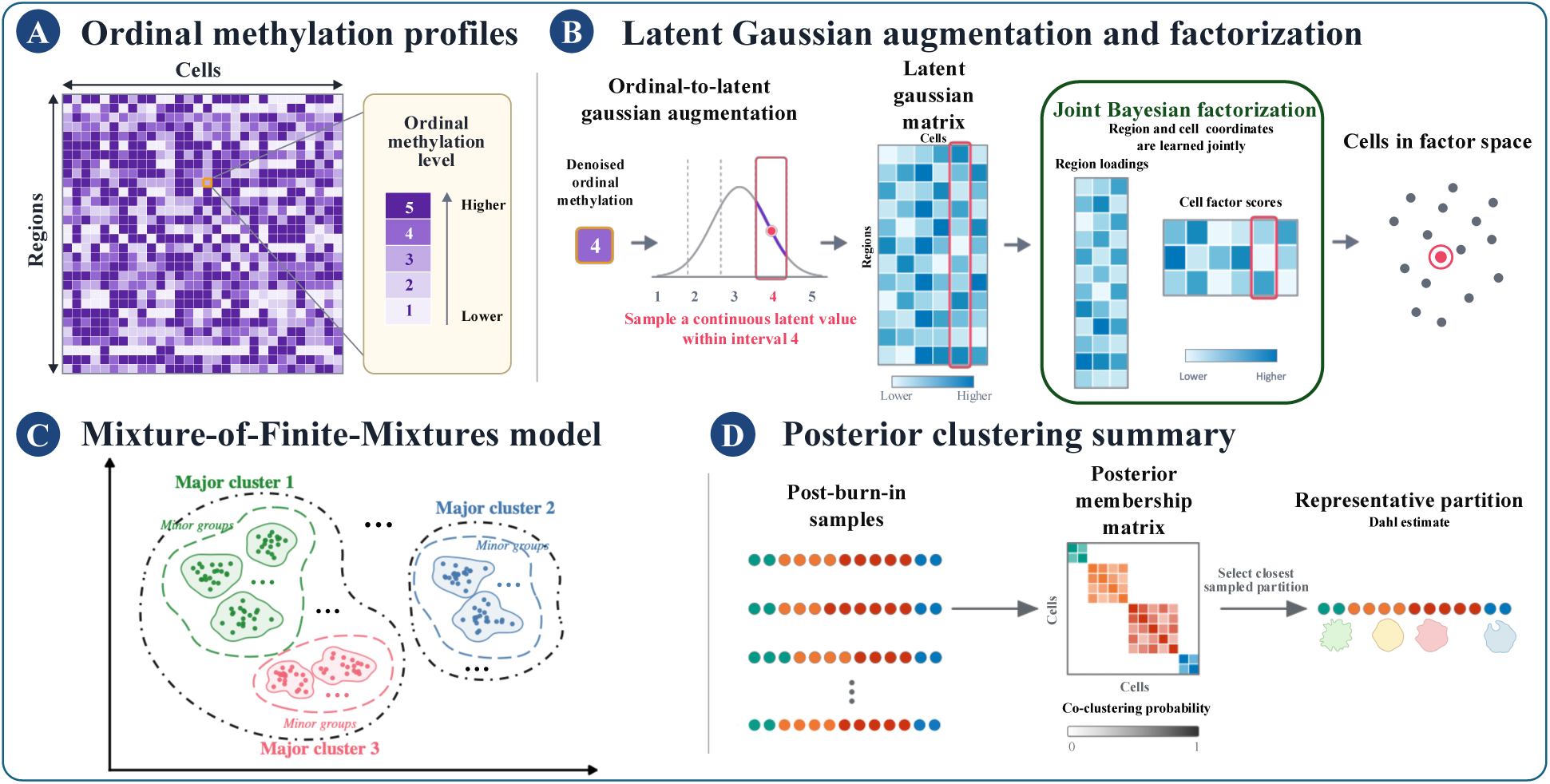
The schematic overview of *SMORE*. **(A)** DNA methylation proportions are converted into ordered methylation states, providing an interpretable ordinal representation of the input data while reducing sensitivity to small measurement fluctuations. **(B)** Latent Gaussian augmentation introduces a continuous variable underlying each ordinal observation, enabling probabilistic modeling of the ordered methylation states. Joint Bayesian factorization then summarizes coordinated methylation variation across genomic regions through feature loadings and represents individual cells by low-dimensional latent factor scores. **(C)** The latent factor scores are modeled using a Mixture-of-Finite-Mixtures (MFM) prior to jointly infer cell-population assignments and the number of occupied clusters, without requiring the number of clusters to be prespecified. Solid, dashed, and dot-dashed contours illustrate cluster-specific covariance structures for candidate partitions, respectively. Diamonds, triangles, and squares denote the corresponding cluster centers. **(D)** Posterior samples are summarized using a co-clustering probability matrix, which quantifies uncertainty in pairwise cell assignments, and a representative partition selected to provide a single interpretable clustering solution.

For each ordinal methylation state, *SMORE* introduces a latent Gaussian variable whose position relative to a set of ordered cutpoints determine the observed methylation state (Fig. 1B). A low-rank factorization of these latent variables captures coordinated methylation patterns across genomic regions: the loading matrix describes region-level contributions, while the factor scores provide low-dimensional coordinates for individual cells. An Mixture-of-Finite-Mixtures (MFM) prior is placed on the cell factor scores, partitioning cells in the latent space while jointly inferring the number of occupied populations (Fig. 1C). This construction avoids fitting mixture components directly to the original high-dimensional profiles and does not require the number of clusters to be fixed before model fitting.

Posterior inference is performed using a data-augmentation Gibbs sampler that updates the latent variables, cutpoints, factor loadings, cell factor scores, cluster assignments and cluster-specific parameters. The retained samples are summarized by a posterior co-clustering probability matrix, whose entries quantify the probability that pairs of cells belong to the same population, and by a representative partition selected using a least-squares criterion (Fig. 1D). Subsequent analyses assess recovery of the cell partition primarily using the Adjusted Rand index(ARI), with normalized mutual information (NMI), adjusted mutual information (AMI), homogeneity and completeness as complementary measures. Detailed model specification and computation are provided in Section 4; metric definitions, hyperparameter sensitivity analyses to the latent dimension *q* and runtime summaries for the distribution-mixture simulations and real-data analyses are reported in the Supplementary Materials Sections S5, S8.5, S7.2 and S8.6, respectively.

### 2.2 Model based simulations

*SMORE* was first evaluated using *in-silico* simulations under an ordinal latent factor mixture model, for which both the underlying cell partition and the number of cell populations were known. The true number of clusters was fixed at *K*_0_ = 5, with *p* = 5,000 genomic features and a true latent dimension of *q*_0_ = 4. Four sample sizes, *n ∈ {*500, 1,000, 2,000, 4,000*}*, were examined under a factorial design combining balanced or unbalanced cluster sizes with strong or weak signal, yielding 16 simulation settings. The strong-and weak-signal settings differed in the degree of cluster separation, the proportion of informative features, and the magnitude of within-cluster variation, with the weak-signal setting producing greater overlap among cell populations. The exact parameter values and the constructions of parameters are provided in the Supplementary Materials Section S6. Cluster sizes were equal in the balanced settings, whereas the unbalanced settings used cluster proportions of (0.40, 0.30, 0.15, 0.10, 0.05). Twenty independent datasets were generated for each scenario.

Briefly, cell-specific latent factor scores were sampled from a five-component Gaussian mixture in the latent space, with cluster-specific covariance structures that allowed differences in shape and volume. Latent Gaussian measurements were then generated through a low-rank factor model in which only a prespecified subset of genomic features had nonzero factor loadings. These measurements were discretized into five ordinal methylation levels using empirical cutpoints chosen to yield marginal category proportions of ***π***^ord^ = (0.15, 0.10, 0.15, 0.20, 0.40). This model-aligned construction generated observations of the same ordinal form analyzed by *SMORE* while retaining a known latent cell-population structure. Full parameter settings and data-generation details are provided in the Supplementary Materials Section S6.

As a complementary evaluation, we conducted an empirically calibrated distribution-mixture simulation based on feature-level methylation distributions estimated from an independent mouse pituitary single-nucleus DNA methylation dataset. Simulated features followed mixtures of two Beta components and point masses at zero and one, with cluster-specific mixture weights controlling population separation. This data-driven design preserved zero/one inflation and distributional heterogeneity observed in real methylation data without generating observations directly from the ordinal latent factor mixture model; full details and results are provided in the Supplementary Materials Section S7.

**Table 1:** Model-based simulation performance of *SMORE* and the ten highest-ranked benchmark pipelines across simulation scenarios. For each scenario, ARI and label-matched clustering accuracy are reported. Values for *SMORE* are shown in boldface, and the best-performing existing method in each column is underlined.

| Method | Balanced strong |  |  |  |  |  |  |  | Unbalanced strong |  |  |  |  |  |  |  |
| --- | --- | --- | --- | --- | --- | --- | --- | --- | --- | --- | --- | --- | --- | --- | --- | --- |
| | $n = 500$ | | $n = 1000$ | | $n = 2000$ | | $n = 4000$ | | $n = 500$ | | $n = 1000$ | | $n = 2000$ | | $n = 4000$ | |
|  | ARI | Acc. | ARI | Acc. | ARI | Acc. | ARI | Acc. | ARI | Acc. | ARI | Acc. | ARI | Acc. | ARI | Acc. |
| **SMORE** | **0.987** | **0.995** | **0.988** | **0.995** | **0.982** | **0.993** | **0.984** | **0.993** | **0.979** | **0.979** | **0.989** | **0.995** | **0.985** | **0.994** | **0.990** | **0.996** |
| UMAP + DBSCAN | 0.786 | 0.774 | <u>0.842</u> | <u>0.824</u> | 0.754 | 0.775 | 0.753 | 0.749 | 0.774 | 0.762 | <u>0.856</u> | <u>0.854</u> | <u>0.758</u> | <u>0.801</u> | <u>0.850</u> | <u>0.814</u> |
| PCA + Hierarchical | 0.747 | 0.763 | 0.816 | 0.807 | <u>0.814</u> | <u>0.844</u> | <u>0.806</u> | <u>0.819</u> | 0.698 | 0.766 | 0.653 | 0.721 | 0.662 | 0.740 | 0.713 | 0.748 |
| UMAP + Hierarchical | 0.792 | 0.791 | 0.795 | 0.774 | 0.762 | 0.778 | 0.779 | 0.768 | 0.598 | 0.670 | 0.591 | 0.648 | 0.625 | 0.697 | 0.629 | 0.686 |
| t-SNE + Hierarchical | 0.800 | 0.790 | 0.827 | 0.800 | 0.732 | 0.739 | 0.553 | 0.598 | 0.575 | 0.659 | 0.457 | 0.601 | 0.444 | 0.605 | 0.513 | 0.669 |
| NMF + Hierarchical | 0.782 | 0.771 | 0.773 | 0.764 | 0.769 | 0.789 | 0.760 | 0.754 | 0.690 | 0.732 | 0.648 | 0.713 | 0.652 | 0.701 | 0.736 | 0.746 |
| UMAP + Kmeans | 0.707 | 0.688 | 0.772 | 0.750 | 0.753 | 0.752 | 0.772 | 0.749 | 0.666 | 0.693 | 0.559 | 0.620 | 0.631 | 0.701 | 0.620 | 0.663 |
| NMF + DBSCAN | 0.763 | 0.773 | 0.757 | 0.771 | 0.701 | 0.749 | 0.530 | 0.571 | <u>0.828</u> | <u>0.872</u> | 0.664 | 0.762 | 0.528 | 0.691 | 0.557 | 0.649 |
| PCA + DBSCAN | 0.816 | 0.843 | 0.666 | 0.707 | 0.612 | 0.708 | 0.622 | 0.653 | <u>0.778</u> | <u>0.842</u> | 0.657 | 0.769 | 0.607 | 0.761 | 0.635 | 0.706 |
| t-SNE + GMM | <u>0.972</u> | <u>0.988</u> | 0.306 | 0.423 | 0.163 | 0.333 | 0.145 | 0.319 | 0.763 | 0.800 | 0.660 | 0.749 | 0.335 | 0.554 | 0.298 | 0.547 |
| UMAP + GMM | 0.752 | 0.743 | 0.807 | 0.776 | 0.786 | 0.807 | 0.730 | 0.740 | 0.607 | 0.677 | 0.650 | 0.701 | 0.554 | 0.602 | 0.582 | 0.633 |

| Method | Balanced weak |  |  |  |  |  |  |  | Unbalanced weak |  |  |  |  |  |  |  |
| --- | --- | --- | --- | --- | --- | --- | --- | --- | --- | --- | --- | --- | --- | --- | --- | --- |
| | $n = 500$ | | $n = 1000$ | | $n = 2000$ | | $n = 4000$ | | $n = 500$ | | $n = 1000$ | | $n = 2000$ | | $n = 4000$ | |
|  | ARI | Acc. | ARI | Acc. | ARI | Acc. | ARI | Acc. | ARI | Acc. | ARI | Acc. | ARI | Acc. | ARI | Acc. |
| **SMORE** | **0.909** | **0.931** | **0.976** | **0.990** | **0.952** | **0.980** | **0.955** | **0.981** | **0.916** | **0.944** | **0.967** | **0.976** | **0.972** | **0.987** | **0.956** | **0.979** |
| UMAP + DBSCAN | <u>0.709</u> | <u>0.771</u> | <u>0.832</u> | <u>0.845</u> | <u>0.784</u> | <u>0.817</u> | 0.647 | 0.672 | <u>0.727</u> | <u>0.816</u> | <u>0.802</u> | <u>0.833</u> | <u>0.799</u> | <u>0.822</u> | <u>0.756</u> | <u>0.781</u> |
| PCA + Hierarchical | 0.638 | 0.711 | 0.695 | 0.720 | 0.689 | 0.733 | 0.641 | 0.711 | 0.598 | 0.660 | 0.610 | 0.688 | 0.647 | 0.733 | 0.601 | 0.675 |
| UMAP + Hierarchical | 0.655 | 0.712 | 0.742 | 0.744 | 0.718 | 0.733 | 0.711 | <u>0.743</u> | 0.548 | 0.638 | 0.575 | 0.656 | 0.582 | 0.684 | 0.591 | 0.664 |
| t-SNE + Hierarchical | 0.676 | 0.708 | 0.565 | 0.623 | 0.707 | 0.726 | 0.309 | 0.429 | 0.495 | 0.640 | 0.420 | 0.571 | 0.611 | 0.683 | 0.466 | 0.622 |
| NMF + Hierarchical | 0.627 | 0.683 | 0.687 | 0.707 | 0.649 | 0.680 | 0.610 | 0.670 | 0.574 | 0.643 | 0.609 | 0.685 | 0.553 | 0.643 | 0.607 | 0.675 |
| UMAP + Kmeans | 0.664 | 0.716 | 0.716 | 0.710 | 0.721 | 0.724 | 0.692 | 0.709 | 0.440 | 0.569 | 0.579 | 0.642 | 0.499 | 0.599 | 0.504 | 0.584 |
| NMF + DBSCAN | 0.390 | 0.477 | 0.652 | 0.718 | 0.419 | 0.518 | 0.225 | 0.323 | 0.590 | 0.660 | 0.712 | 0.745 | 0.553 | 0.672 | 0.347 | 0.518 |
| PCA + DBSCAN | 0.377 | 0.509 | 0.458 | 0.552 | 0.424 | 0.546 | 0.194 | 0.344 | 0.433 | 0.642 | 0.606 | 0.726 | 0.621 | 0.735 | 0.495 | 0.612 |
| t-SNE + GMM | 0.462 | 0.607 | 0.320 | 0.457 | 0.306 | 0.413 | 0.159 | 0.314 | 0.486 | 0.650 | 0.367 | 0.572 | 0.652 | 0.753 | 0.472 | 0.651 |
| UMAP + GMM | 0.639 | 0.681 | 0.768 | 0.778 | 0.717 | 0.726 | <u>0.712</u> | 0.738 | 0.531 | 0.637 | 0.582 | 0.673 | 0.594 | 0.689 | 0.549 | 0.629 |

## Results

*SMORE* accurately recovered the true latent partition across all model-based simulation scenarios. Recovery was nearly perfect under strong signal: mean ARI values were 0.987, 0.988, 0.982, and 0.984 for *n* = 500, 1,000, 2,000, and 4,000, respectively; with unbalanced cluster sizes, mean ARI ranged from 0.979 to 0.990. At *n* = 1,000, *SMORE* ranked first across the balanced–rstrong, balanced–weak, unbalanced–strong, and unbalanced–weak scenarios, demonstrating consistently strong recovery across changes in signal strength and cluster-size balance (Fig. 2A). This advantage was also consistent across ARI, NMI, AMI, homogeneity, and completeness, indicating that the inferred partitions captured both agreement with the true cell assignments and the overall population structure (Fig. 2B).

**Figure 2:**
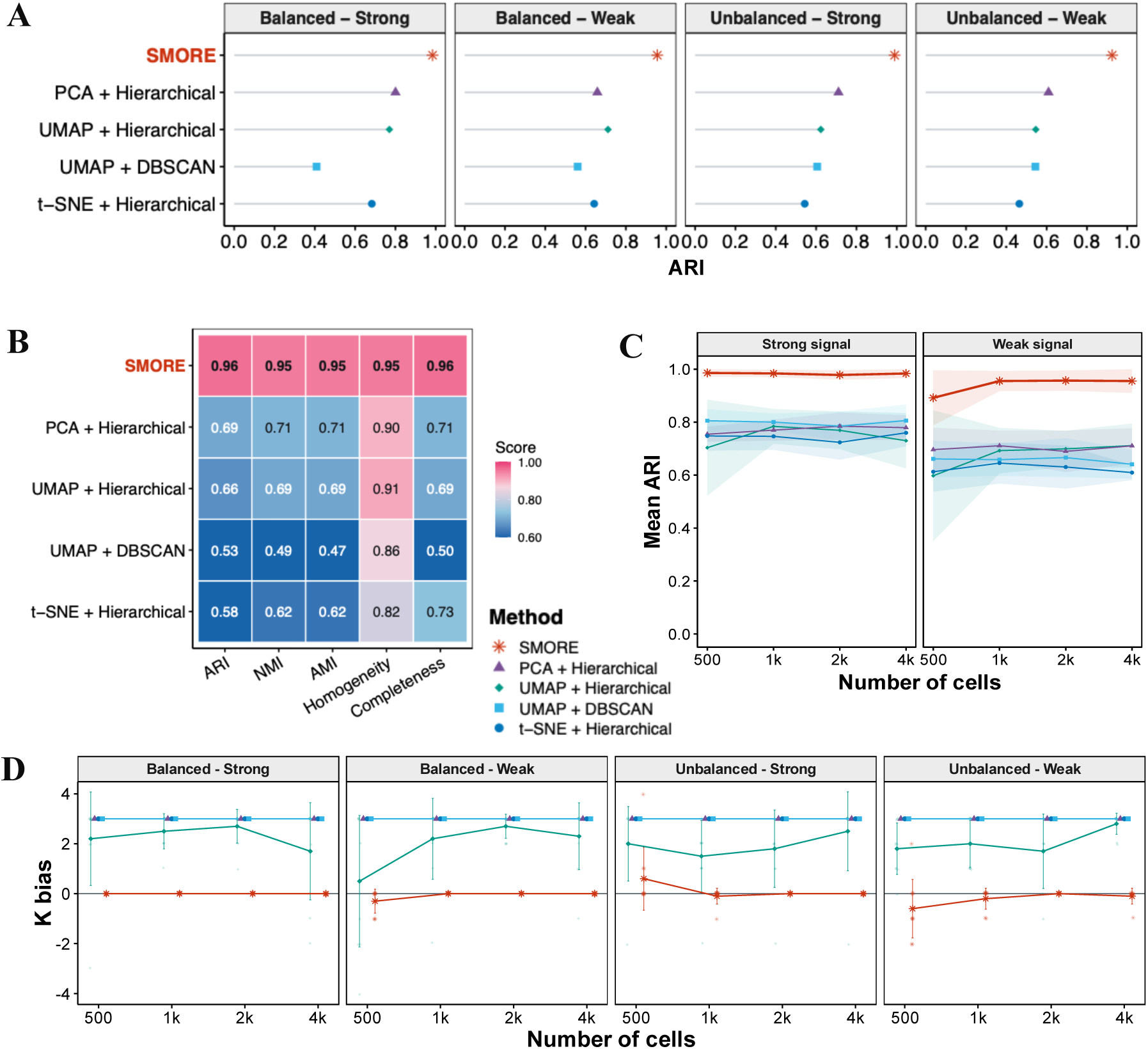
Performance comparison under the model-based simulation settings. **(A)** ARI-based method rankings at *n* = 1,000 cells under four scenarios: balanced–strong, balanced–weak, unbalanced–strong, and unbalanced–weak. **(B)** Performance in terms of ARI, NMI, AMI, homogeneity, and completeness at *n* = 1,000 cells, averaged across the four simulation scenarios. **(C)** Scalability under the balanced scenarios as the sample size increases. Lines indicate mean ARI, and shaded ribbons indicate *±* one standard deviation across replicates. **(D)** Bias in the estimated number of clusters across sample sizes and simulation scenarios. Small translucent points represent individual replicates, large symbols represent replicate means, and error bars indicate *±* one standard deviation. The horizontal line at zero indicates no bias relative to the true number of clusters, *K*_0_ = 5.

Recovery remained high when weaker signal produced greater overlap between populations. In the balanced settings, mean ARI values were 0.909, 0.976, 0.952, and 0.955 for *n* = 500, 1,000, 2,000, and 4,000, respectively. With unbalanced cluster sizes, mean ARI increased from 0.916 at *n* = 500 to 0.967 at *n* = 1,000 and 0.972 at *n* = 2,000, and remained high at 0.956 for *n* = 4,000. Increasing the number of cells produced the largest gains in the more difficult weak-signal settings, although performance did not improve monotonically at every sample size (Fig. 2C). Notably, *SMORE* achieved the highest ARI among the evaluated methods even at the smallest sample size of *n* = 500, indicating that its advantage was retained when relatively few cells were available.

Among the benchmark methods, PCA combined with hierarchical clustering was the most consistently competitive, achieving the highest ARI among the benchmark pipelines across the four scenarios at *n* = 1,000. NMF or UMAP followed by hierarchical clustering, as well as UMAP followed by a Gaussian mixture model (GMM), performed well in selected settings but were less stable across changes in signal strength and cluster-size balance. Several benchmark pipelines deteriorated markedly under weak signal. In contrast, *SMORE* maintained estimated-cluster-number bias close to zero across sample sizes and scenarios, whereas the benchmark pipelines showed larger and more variable biases (Fig. 2D).

The detailed cluster assignment showed that *SMORE* more clearly recovered the underlying cluster structure, particularly at larger sample sizes (Fig. 3). The Sankey flows further showed an almost one-to-one correspondence between the true and inferred clusters for *SMORE* at *n* = 1,000 and *n* = 2,000. Whereas several benchmark methods continued to split indivisual true clusters or merge distinct groups; notably, tSne + GMM assign nearly all cells to a single inferred cluster at *n* = 2,000. These simulations above establish the recovery performance of *SMORE* when data follow an ordinal latent factor mixture structure.

**Figure 3:**
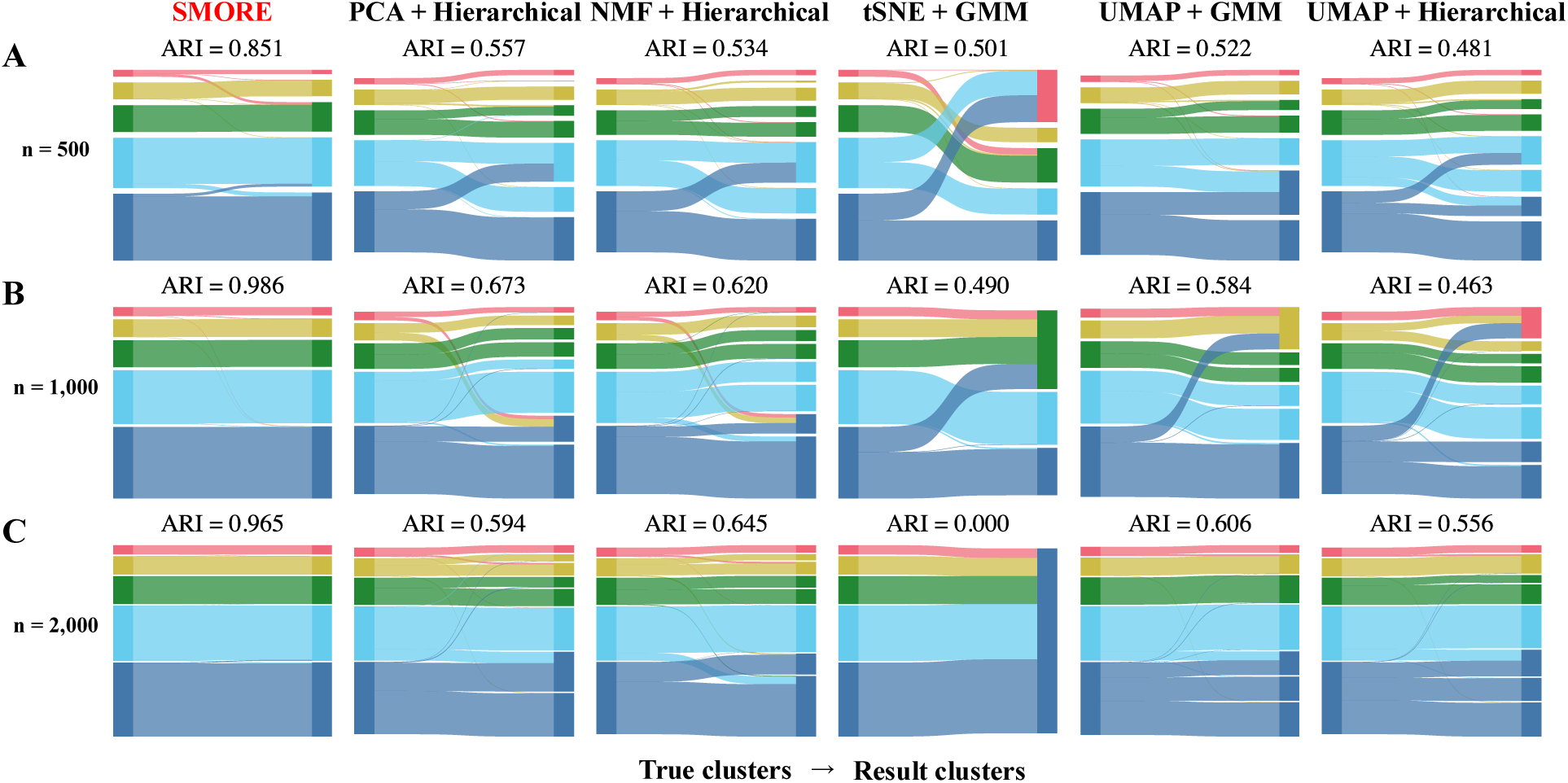
Comparison of clustering assignment accuracies. Sankey diagrams show the correspondence between the true clusters and those inferred by *SMORE* and five existing methods in simulation replicates at **(A)** *n* = 500, **(B)** *n* = 1,000, and **(C)** *n* = 2,000. The width of each flow is proportional to the number of cells shared by the corresponding true and inferred clusters.

### 2.3 Real data analysis

We evaluated *SMORE* on two human single-cell DNA methylation datasets representing distinct tissue and cellular contexts. The lung dataset comprised ten major cell-type groups, the PBMC dataset comprised five major peripheral blood cell-type groups, and the primary motor cortex dataset comprised ten major cell-type groups. For each dataset, methylation proportions were calculated from the methylated-read counts and total coverage for each 5-kb genomic region, as in the original publication (Zhou et al., 2026). Cells and genomic regions with excessive missingness were removed. Complete filtering criteria and implementation details are provided in the Supplementary Materials Section S8.1.

The resulting methylation levels were discretized into five ordered categories defined by equally spaced cutpoints on [0, 1]. This ordinal representation preserves the ordering from low to medium to high methylation status while reducing sensitivity to small fluctuations in the estimated methylation proportions. For each dataset, the same processed ordinal matrix was used as input for the all models.

After preprocessing, the lung dataset contained 507 cells and 15,223 retained 5-kb genomic regions, while the PBMC dataset contained 1,975 cells and 15,239 retained regions and the primary motor cortex dataset contained 540 cells and 15,430 retained regions. Available cell-type annotations were obtained from the original publication as reference labels for evaluation.

## Results

We benchmarked *SMORE* against 25 existing pipelines formed by combining five dimension-reduction methods with five clustering algorithms. Complete numerical results for *SMORE* and the ten highest-performing benchmark pipelines are reported in Table 2. *SMORE* achieved the highest ARI (0.822), followed by t-SNE with K-means (ARI = 0.807), t-SNE with hierarchical clustering (ARI = 0.800), and ICA with a Gaussian mixture model (ARI = 0.749). Figure 4A summarizes the ARI-based ranking of *SMORE* and the four highest-performing benchmark pipelines. Although several benchmark pipelines recovered substantial reference structure, their performance remained lower than that of *SMORE* and varied considerably across combinations of dimension-reduction and clustering methods.

**Figure 4:**
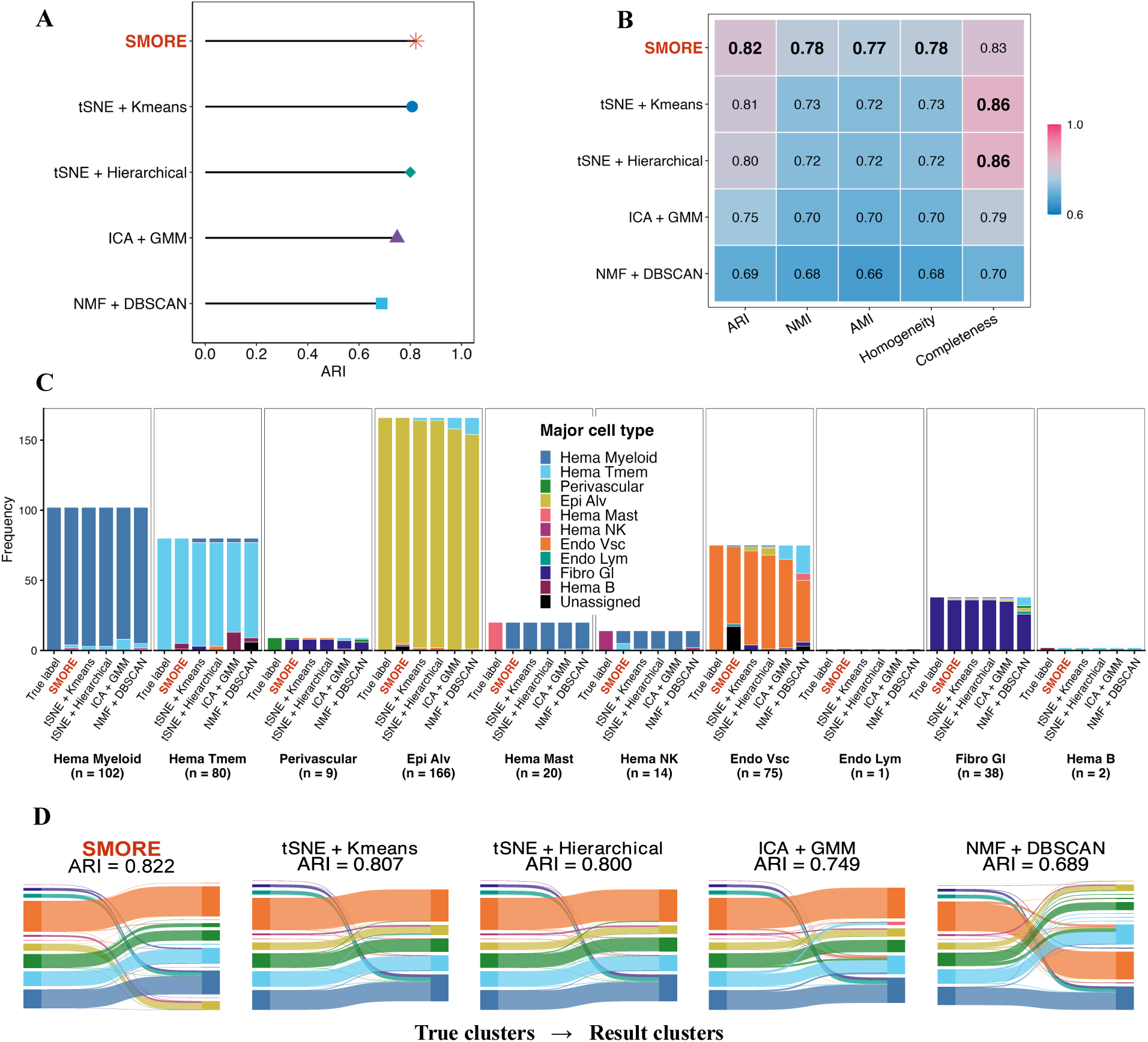
Benchmarking clustering of real single-cell DNA methylation dataset from lung. **(A)** Ranking of *SMORE* and the four highest-performing benchmark pipelines by ARI. **(B)** Comparison of ARI, NMI, AMI, homogeneity, and completeness. **(C)** Composition of the clusters inferred by each method across the ten reference cell groups. Inferred clusters were matched to reference labels based on maximum overlap, and clusters without a unique match are shown as unassigned. **(D)** Sankey diagrams showing the correspondence between the reference labels and the clusters inferred by *SMORE* and the four highest-performing benchmark pipelines.

**Table 2:** Clustering performance of *SMORE* and the ten highest-performing existing methods on the lung dataset (*n* = 507 cells). Values for *SMORE* are shown in boldface, and the best-performing benchmark value in each column is underlined.

| Method | ARI | NMI | AMI | Hom. | Comp. |
| --- | --- | --- | --- | --- | --- |
| **SMORE** | **0.822** | **0.782** | **0.774** | **0.782** | **0.825** |
| t-SNE + K-means | <u>0.807</u> | <u>0.727</u> | <u>0.721</u> | 0.727 | <u>0.856</u> |
| t-SNE + Hierarchical | 0.800 | 0.722 | 0.717 | 0.722 | 0.856 |
| ICA + GMM | 0.749 | 0.704 | 0.696 | 0.704 | 0.785 |
| NMF + DBSCAN | 0.689 | 0.675 | 0.659 | 0.675 | 0.700 |
| PCA + DBSCAN | 0.668 | 0.649 | 0.634 | 0.649 | 0.677 |
| t-SNE + DBSCAN | 0.667 | 0.526 | 0.482 | 0.733 | 0.526 |
| PCA + K-means | 0.633 | 0.578 | 0.571 | 0.578 | 0.798 |
| ICA + Louvain | 0.628 | 0.693 | 0.684 | 0.733 | 0.693 |
| NMF + GMM | 0.626 | 0.614 | 0.605 | 0.614 | 0.715 |
| NMF + Louvain | 0.606 | 0.654 | 0.643 | <u>0.740</u> | 0.654 |

The advantage of *SMORE* was also evident in NMI, AMI, and homogeneity (Fig. 4B). Its NMI and AMI were 0.782 and 0.774, respectively, compared with 0.727 and 0.721 for t-SNE followed by K-means. Completeness was the only reported metric for which *SMORE* did not rank first (0.825 versus 0.856 for both leading t-SNE pipelines), indicating a trade-off between its stronger overall agreement and homogeneity and the slightly higher completeness of the t-SNE solutions.

Cluster composition and Sankey diagram revealed differences in partition granularity (Fig. 4C, D). *SMORE* inferred 11 clusters, close to the 10 reference cell groups, whereas each of the two leading t-SNE pipelines inferred five clusters. All three approaches recovered the major lung cell groups, but the five-cluster solutions mistakenly merged several smaller groups. At the cell-type level, *SMORE* showed clear correspondence for the abundant alveolar epithelial, hematopoietic myeloid, hematopoietic memory T-cell, and gastrointestinal fibroblast groups. In contrast, the smaller hematopoietic mast-cell, hematopoietic natural killer-cell, perivascular, lymphatic endothelial, and hematopoietic B-cell groups generally did not form distinct inferred clusters and were instead absorbed into clusters dominated by larger populations. The vascular endothelial group was partially subdivided across two inferred clusters. Overall, the 11-cluster solution primarily reflected subdivisions within major cell-type groups rather than uniform recovery of every annotated cell group.

Multidimensional scaling (MDS) provided a cell-level view of the clustering assignments in the lung dataset (Figure 5A). Gray points denote assignments to the primary matched cluster, while blue points indicate within-group over-splitting and magenta points indicate assignment to a cluster dominated by a different reference group. *SMORE* assigned most cells to their primary matched clusters; 24 cells were assigned to additional clusters within the same reference cell group, and 57 were assigned to clusters dominated by a different reference group. In contrast, PCA followed by Louvain clustering assigned 91 cells to additional clusters within the same reference cell group and 71 cells to clusters dominated by a different reference group, resulting in twice as many cells outside their primary matched clusters as *SMORE*. Among the representative PCA-based pipelines, PCA followed by K-means achieved the highest ARI of 0.729, inferred eight clusters, and assigned 76 cells to clusters dominated by a different reference group. PCA followed by DBSCAN, Louvain clustering, and GMM achieved lower ARI values of 0.668, 0.619, and 0.594, respectively. The higher agreement achieved by *SMORE* was therefore accompanied by fewer cross-group assignments and greater preservation of within-group structure.

**Figure 5:**
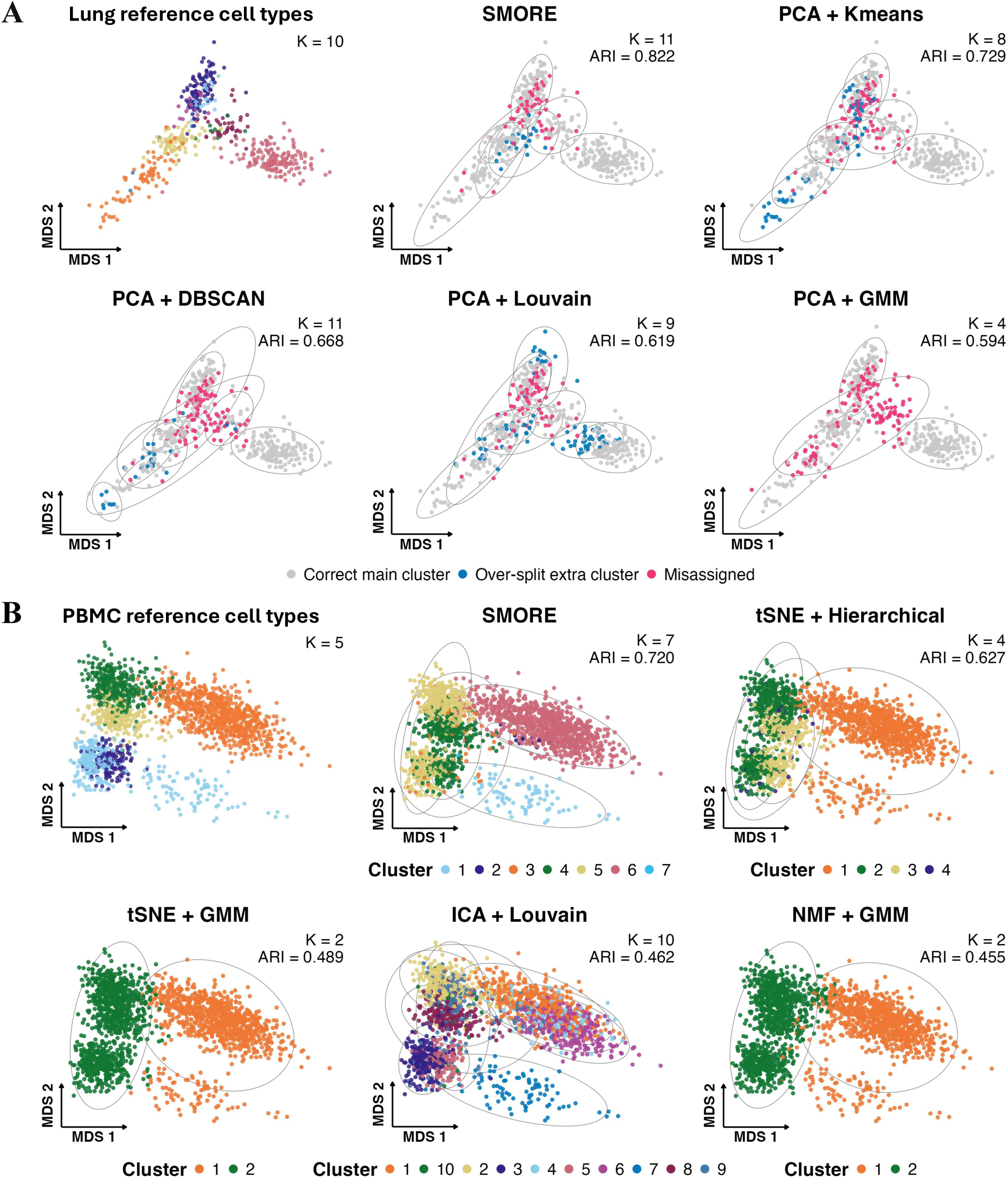
Multidimensional scaling visualization in **(A)** Lung and **(B)** PBMC. Within each dataset, all panels use the same MDS coordinates calculated from the common processed methylation matrix, allowing clustering assignments to be compared without changing the underlying visualization. The leftmost panel shows the reference cell-group labels. In **(A)**, gray points denote cells assigned to the primary cluster associated with their reference group, blue points denote cells assigned to additional clusters associated with the same reference group and therefore indicate within-group over-splitting, and pink points denote cells assigned to clusters dominated by a different reference group. Ellipses outline the clusters inferred. In **(B)**, colors denote inferred cluster memberships and are specific to each method. ^17^

Similarly, we evaluated the performance on the PBMC dataset, which comprised 1,975 cells from five reference cell types (Figure 5B). *SMORE* achieved an ARI of 0.720 and inferred seven clusters. By comparison, t-SNE followed by hierarchical clustering achieved an ARI of 0.627 and inferred four clusters, whereas t-SNE followed by GMM achieved an ARI of 0.489 and inferred two clusters. Independent component analysis followed by Louvain clustering achieved an ARI of 0.462 and inferred 10 clusters, whereas non-negative matrix factorization followed by GMM achieved an ARI of 0.455 and inferred two clusters. These benchmark solutions therefore tended either to merge reference groups or to divide them into multiple clusters. Such variation likely reflects the two-stage design of these benchmark pipelines, in which dimension reduction and clustering are optimized separately, as well as their dependence on method-specific distributional assumptions, graph construction, and cluster-selection criteria. In contrast, *SMORE* models methylation measurements as ordinal states and jointly estimates the latent representation, cell assignments, and number of clusters, helping it avoid these more extreme partitions and achieve higher agreement with the reference labels. This greater stability was broadly consistent with the smaller and less variable cluster-number bias observed for *SMORE* in the simulations.

Across the lung and PBMC results presented in the main text, *SMORE* achieved greater agreement with the reference labels than the benchmark pipelines shown in the main text while preserving a more granular representation of cell-group structure. Additional analyses for all three real datasets are provided in Supplementary Materials Sections S8.2–S8.6.

## 3 Discussion

We developed *SMORE*, a Bayesian framework that integrates ordinal measurement modeling, dimension reduction, and clustering for single-cell DNA methylation data. The ordinal observation model preserves the ordering of methylation levels while reducing sensitivity to small fluctuations arising from sparse and heterogeneous coverage. Jointly modeling the latent factors and their MFM-based partition allows the latent representation and population structure to inform each other, while supporting inference of an unknown finite number of populations with unequal abundances and discouraging spurious small clusters. Across model-based simulations, *SMORE* accurately recovered the underlying cell partitions and population numbers, including under weak signal and unequal cluster sizes. Across the three real datasets, it recovered biologically supported structure, although its relative performance depended on cell-population composition and the granularity of the reference annotations. It achieved the highest agreement in the lung and peripheral blood datasets, whereas the primary motor cortex analysis illustrated a trade-off between retaining finer population structure and maximizing an ARI dominated by abundant cell groups. Beyond these comparisons, *SMORE* provides posterior estimates of the population number and cell co-clustering probabilities, allowing uncertainty in the inferred partition to be assessed directly.

The relative advantage of *SMORE* was most apparent when populations were weakly separated or cluster sizes were unbalanced, which aligns well with the intrinsic characteristics of single-cell DNA methylation data. Jointly estimating the latent representation and population structure makes the analysis less sensitive to the choice of separate dimension-reduction and clustering procedures. The value of *SMORE* therefore lies in the combination of clustering performance and inferential output, particularly when the number of populations or the assignment of individual cells is uncertain.

It is worth noting that the form of the reference labels differed across datasets. For the lung dataset, we used the annotated cell-type labels from the source study, whereas for the PBMC and primary motor cortex datasets, we retained the deposited cluster identifiers without mapping them to the available biological names. These labels were derived from DNA methylation and 3D chromatin-contact profiles, with biological annotation further informed by scRNA-seq and snATAC-seq data (Zhou et al., 2026). Because they were analytically inferred and partly based on DNA methylation, they are best regarded as a silver standard rather than an independent gold standard. Accordingly, the reported metrics quantify agreement with the published reference structure rather than definitive cell-type identification accuracy. Finer subdivisions identified by *SMORE* require independent biological validation.

Several limitations should also be considered. When the signal is strong and populations are well separated, conventional two-stage workflows may achieve comparable clustering performance at substantially lower computational cost. The joint Bayesian inference and uncertainty quantification provided by *SMORE* require longer runtimes, which may limit scalability to larger datasets. Discretizing methylation proportions also reduces sensitivity to small continuous fluctuations but removes variation within each ordinal category, while fixed cut-off points may not capture feature-specific measurement characteristics. In addition, the current workflow relies on filtering and imputation rather than modeling sequencing coverage and missingness directly; count-level observation models could propagate coverage uncertainty more fully. Finally, *SMORE* assumes that methylation variation follows a low-dimensional linear factor structure and that cell populations form a finite mixture in this space. Although these assumptions provide a tractable framework, they may be less appropriate for strongly nonlinear or continuous population structures, such as developmental trajectories. Improving computational efficiency will be important for extending the framework to larger datasets.

## 4 Methods

### 4.1 Bayesian ordinal factor mixture-of-finite-mixtures model

#### Ordinal Factor Model with Latent Gaussian Augmentation

We develop a Bayesian framework for joint dimension reduction and cell-population discovery from single-cell methylation profiles. The model integrates three components: an ordinal representation of methylation measurements, a low-dimensional latent factor model, and a mixture-of-finite-mixtures (MFM) model for cell clustering. By estimating these components jointly, uncertainty in the observed methylation profiles is propagated through the latent representation and into the inferred cell populations. Let **Y** = (*Y_ij_*) *∈ {*1*,…, C}^p×n^* denote the ordinal methylation matrix after preprocessing, where *Y_ij_* represents the methylation level of genomic region *i* = 1*,…, p* in cell *j* = 1*,…, n*. Here, *C* denotes the total number of ordered methylation categories; in our analyses, *C* = 5, corresponding to five levels ordered from the lowest to the highest methylation proportion. Modeling methylation measurements on an ordinal scale preserves their natural ordering from low to high methylation levels, while reducing sensitivity to small fluctuations in estimated methylation proportions caused by sparse and heterogeneous sequencing coverage. To model the ordinal observations, a latent continuous variable *Z_ij_* is introduced for each *Y_ij_*using latent Gaussian augmentation (Albert & Chib, 1993). Let ***γ*** = (*γ*_0_*,…, γ_C_*) denote a set of ordered cutpoints, with *γ*_0_ = *−∞* and *γ_C_* = *∞*. The observed ordinal level is related to the latent variable through

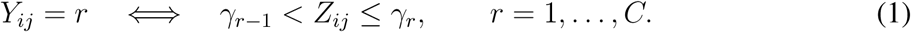

For identifiability of the ordinal probit model, *γ*_1_ = 0 is fixed and the latent error variance is fixed at one. Single-cell methylation profiles are high dimensional, whereas the major biological variation across cells is expected to be characterized by a smaller number of coordinated methylation patterns. We therefore impose a low-rank factor structure on the latent Gaussian variables. Let **Z***_j_* = (*Z*_1_*_j_,…, Z_pj_*)*^⊤^* denote the latent methylation profile for cell *j*, and assume that

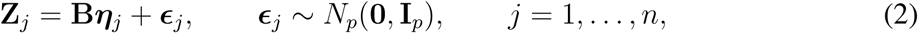

where **B** *∈* R*^p×q^* is the factor loading matrix and ***η****_j_ ∈* R*^q^* is the latent factor score for cell *j*, with *q ≪ p*. Thus, the original high-dimensional ordinal methylation profile is represented by the low-dimensional coordinate ***η****_j_*. Importantly, the latent factor scores are not estimated in a separate dimension-reduction step. Instead, they are inferred jointly with the ordinal latent variables and the cell-population assignments. This joint formulation allows information across genomic regions to contribute simultaneously to estimation of the latent representation and identification of cell populations. The factor representation is invariant to orthogonal rotations. A positive lower-triangular constraint is therefore imposed on the leading *q × q* block of **B** to reduce rotational and sign indeterminacy and provide a stable orientation for posterior computation (Geweke & Zhou, 1996). Because the mixture components introduced below have cluster-specific covariance matrices, the positive lower-triangular constraint is used to provide a consistent orientation of the latent factor representation. Details of the ordinal transformation, positive lower-triangular constraint, and corresponding likelihood are provided in the Supplementary Materials Sections S1.1 and 1.2.

#### Mixture of Finite Mixtures Clustering of Latent Factors

The latent factor scores provide a low-dimensional representation of the cells, but the number of underlying cell populations is unknown. To jointly infer the number of populations and cell memberships, an MFM prior (Miller & Harrison, 2018) is placed directly on {*ηj*}*^n^_j=1_*. Specifically, we assume

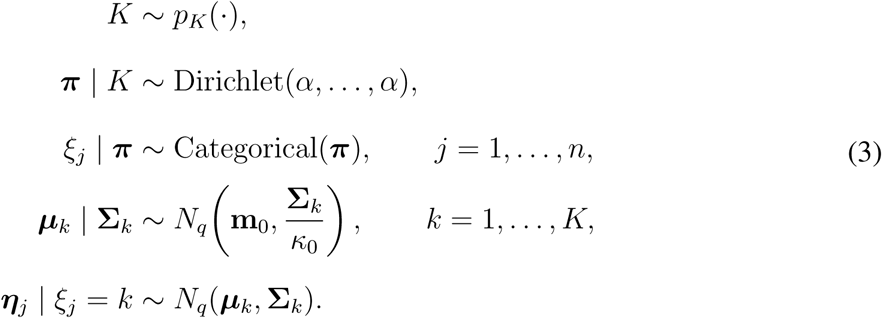

Here *p_K_*is a proper prior on the positive integers, ***π*** contains the mixture weights, and *ξ_j_*denotes the cluster membership of cell *j*. The vector ***µ****_k_* represents the location of population *k* in the latent factor space, whereas **Σ***_k_* characterizes its within-population covariance structure. Thus, distinct cell populations may differ in both their locations and covariance structures within the latent methylation space. This formulation clusters cells through their low-dimensional factor scores rather than directly through the original *p*-dimensional methylation profiles. The factor model borrows information across genomic regions to estimate the latent representation, while the MFM component simultaneously uses the emerging population structure to inform the factor scores. Consequently, dimension reduction and clustering are performed within a single probabilistic model rather than as separate analytical steps.

#### Prior Specification

For the factor loadings, independent Gaussian priors with variance *σ*^2^ are assigned to the free elements of **B**, subject to the positive lower-triangular constraint. A proper prior subject to the ordering constraint is assigned to the free cutpoints *γ*_2_*,…,γ_C−_*_1_. For the mixture components in (3), we set **m**_0_ = **0** and assign

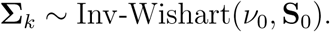

Conditional on **Σ***_k_*, the component-specific mean ***µ****_k_* follows the Gaussian distribution specified in (3). The concentration parameter *α* controls the relative cluster sizes, whereas *p_K_* determines the prior distribution on the number of cell populations. All hyperparameters are fixed before model fitting and are kept unchanged across simulation settings and real-data analyses unless otherwise stated. The numerical values of σ^2^*_B_*, *κ*_0_, *ν*_0_, **S**_0_, *α*, the parameters defining *p_K_*, together with the choice of latent dimension *q*, are provided in Supplementary Materials Section S1.5. *q* controls the flexibility of the low-dimensional representation and was set to *q* = 10 in the primary analyses. To assess sensitivity to this choice, we repeated the real-data analyses using *q ∈ {*5, 10, 15, 20*}* while holding the remaining model settings fixed, and compared clustering performance and computational runtime across these values. Full sensitivity results are provided in Supplementary Materials Section S8.5.

### 4.2 Posterior computation and clustering summary

Posterior inference is performed using an Albert–Chib data-augmentation Gibbs sampler (Albert & Chib, 1993; Neal, 2000). At each iteration, the sampler updates the latent Gaussian variables **Z**, factor scores {**η**_j_}*^n^_j_*_=1_, loading matrix **B**, cutpoints ***γ***, MFM cluster memberships ***ξ***, and cluster-specific parameters {(μ*_k_*,Σ*_k_*)}*_k=1_*^K+^. Conditional on the factor scores and loading matrix, each latent Gaussian variable is sampled from a normal distribution truncated to the interval determined by its observed ordinal methylation level in (1). The factor scores and free elements of the loading matrix have Gaussian full conditional distributions, subject to the positive lower-triangular constraint on **B**. For the clustering step, the mixture weights are integrated out and the allocation variables are updated using the MFM partition probabilities. The resulting collapsed update allows cell assignments and the number of cell populations to be learned directly from the latent factor representation. The complete Gibbs sampling scheme, including all full conditional distributions, MFM allocation probabilities, and their derivations, is provided in the Supplementary Materials Sections S2 and S3. Because mixture labels are arbitrary across posterior samples, clustering uncertainty is summarized using pairwise posterior co-clustering probabilities. For cells *j* and *j^′^*, let

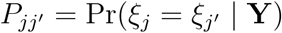

denote the posterior probability that the two cells belong to the same population. The posterior similarity matrix is estimated by averaging the corresponding binary co-clustering matrices across retained MCMC samples. A representative partition is selected using the least-squares partition estimator of Dahl (2006). Specifically, the retained posterior partition whose co-clustering matrix is closest to the posterior similarity matrix in squared distance is selected. This representative partition is used for the clustering results reported in the simulation and real-data analyses, while the posterior similarity matrix provides a direct measure of uncertainty in cell-population assignments. For all analyses, multiple MCMC chains are run from dispersed initial values. Convergence is assessed using trace plots of the log posterior, the inferred number of cell populations, and representative model parameters, together with standard between-chain and within-chain diagnostics. Exact numbers of iterations, burn-in periods, initialization procedures, hyperparameter values, and convergence diagnostics are provided in the Supplementary Materials Sections S1.5 and S9.

### 4.3 Benchmark comparison

Comparator eligibility was defined according to data compatibility and the scope of the controlled evaluation. A benchmark method was eligible if it could operate on the same cell-by-bin matrix used by *SMORE* without requiring unavailable CpG-level observations, genomic coordinates, or reconstruction of within-region methylation profiles. Complete workflows whose principal contributions include upstream feature discovery, coverage normalization, or method-specific data representation were not included because their use would prevent attribution of performance differences specifically to downstream representation and clustering.

For benchmark methods requiring a prespecified number of clusters, the number of clusters was selected using the gap statistic (Tibshirani et al., 2001). ARI was treated as the primary clustering metric, with NMI, AMI, homogeneity, completeness, and label-matched clustering accuracy used as complementary measures. Full tuning procedures and metric definitions are provided in Supplementary Materials Sections S4 and S5.

### 4.4 Data source and preprocessing

Three publicly available human single-cell DNA methylation datasets were analyzed: a lung dataset, a PBMC dataset, and a primary motor cortex dataset. All three datasets were obtained from the recently released atlas of 3D genome organization and DNA methylation in human (Zhou et al., 2026).

For each dataset, the matrices summarize methylated-read counts and total read coverage in non-overlapping 5-kb genomic bins, with rows corresponding to genomic bins and columns to individual cells. The initial lung, PBMC and primary motor cortex dataset matrices comprised 15,933 bins across 573 cells, 15,987 bins across 2,000 cells and 15,954 bins across 542 cells, respectively. Available cell-type annotations from the original publication were retained as reference labels for external evaluation of clustering performance. For each bin and cell, the methylation proportion was calculated by dividing the methylated-read count by the corresponding coverage; entries with zero coverage were treated as missing. Missing values were imputed using a cell-specific kernel density estimation approach, and zero-variance features were excluded. Quality control, cell and feature filtering, missing-value imputation and ordinal transformation are described in Supplementary Materials Section S8.1.

## Data Availability

The three human single-cell DNA methylation datasets analyzed in this study—the lung dataset, the PBMC dataset, and the primary motor cortex dataset—are publicly available through the Human Cell Epigenome Atlas data repository (https://huggingface.co/datasets/zhoujt1994/HumanCellEpigenomeAtlas_sc_allc). The associated publication is Zhou et al. (2026).

An additional mouse pituitary single-nucleus DNA methylation dataset was used only to estimate the feature-level mixture distributions employed in the simulation study. This supporting dataset is publicly available from the NCBI Gene Expression Omnibus under accession GSE152011 (https://www.ncbi.nlm.nih.gov/geo/query/acc.cgi?acc=GSE152011). The associated publication is Ruf-Zamojski et al. (2021).

## Code Availability

*SMORE* is publicly available as an R package at: https://github.com/JaneDeng28/SMORE.git. The repository includes detailed instructions, implementation functions, and reproducible example datasets to guide users through typical workflows. The source code is released under the MIT License.

## Acknowledgments

This work was supported by grants from the National Institutes of Health awarded (NIGMS R35GM154862 to H.F.) and the National Science Foundation (NSF SES-2619578 to G.H. and J.D.). The content is solely the responsibility of the authors and does not necessarily represent the official views of the National Institutes of Health.

## Author Contribution Statement

G.H. and H.F. conceived the study. Z.W. and J.D. performed *in-silico* simulation studies and real data analysis. W.T. acquired and processed real scDNAm datasets. J.D., Z.W., G.H. and H.F. wrote the manuscript. All authors helped edit the final manuscript.

