## Supplementary Materials for "SMORE: joint dimension reduction and cell population discovery on single-cell methylome data"

Jingwen Deng$^{\dagger}$, Zixi Wang$^{\dagger}$, Wen Tang, Guanyu Hu$^{*}$, Hao Feng$^{*}$

† indicates joint first authorship.

∗ indicates corresponding authors.

**S2 Full conditional distributions and their derivations** **11**

**S3 Algorithm** **15**

**S4 Tuning Parameters and Benchmark Settings** **15**
S4.1 Gap statistics . . . . . 15
S4.2 Hyper-parameter . . . . . 17

**S5 Evaluation metrics** **17**

**S6 Model-based simulation design** **18**

**S7 Distribution-mixture simulation** **20**

**S9 Additional analysis of results** **39**

### S1 Additional methodological details of the Bayesian ordinal factor MFM model

#### S1.1 Ordinal likelihood and latent Gaussian representation

The continuous methylation level matrix is converted into an ordinal matrix by discretizing each observed level into one of five ordered categories according to predefined thresholds on $[0, 1]$. Categories are labeled from 1 to 5 in order of increasing methylation level.

Let

$$\mathbf{Y} = (Y_{ij}) \in \{1, \ldots, C\}^{p \times n}$$

denote the ordinal methylation matrix, where $i = 1, \ldots, p$ indexes genomic regions and $j = 1, \ldots, n$ indexes cells. In the applications considered in this study, $C = 5$.

For each observed ordinal response $Y_{ij}$, we introduce a latent continuous variable $Z_{ij}$ and an ordered set of cutpoints

$$-\infty = \gamma_0 < \gamma_1 < \cdots < \gamma_{C-1} < \gamma_C = +\infty,$$

such that

$$Y_{ij} = r \quad \Longleftrightarrow \quad \gamma_{r-1} < Z_{ij} \leq \gamma_r, \qquad r = 1, \ldots, C.$$

Conditional on the latent factor score $\boldsymbol{\eta}_j$, the latent variable follows

$$Z_{ij} \mid \mathbf{B}_i, \boldsymbol{\eta}_j \sim N\left(\mathbf{B}_i^\top \boldsymbol{\eta}_j, 1\right),$$

where $\mathbf{B}_i^\top$ is the $i$th row of the loading matrix $\mathbf{B}$. Equivalently,

$$\mathbf{Z}_j = \mathbf{B}\boldsymbol{\eta}_j + \boldsymbol{\epsilon}_j, \qquad \boldsymbol{\epsilon}_j \sim N_p(\mathbf{0}, \mathbf{I}_p).$$

Integrating out $Z_{ij}$ gives the observed-data category probability

$$\Pr(Y_{ij} = r \mid \mathbf{B}_i, \boldsymbol{\eta}_j, \boldsymbol{\gamma}) = \Pr\left(\gamma_{r-1} < Z_{ij} \leq \gamma_r \mid \mathbf{B}_i, \boldsymbol{\eta}_j\right) = \Phi\left(\gamma_r - \mathbf{B}_i^\top \boldsymbol{\eta}_j\right) - \Phi\left(\gamma_{r-1} - \mathbf{B}_i^\top \boldsymbol{\eta}_j\right),$$

where $\Phi(\cdot)$ denotes the standard normal distribution function. Therefore, the observed-data likelihood is

$$L(\mathbf{B}, \boldsymbol{\eta}, \boldsymbol{\gamma}; \mathbf{Y}) = \prod_{j=1}^{n} \prod_{i=1}^{p} \left[ \Phi\left(\gamma_{Y_{ij}} - \mathbf{B}_i^\top \boldsymbol{\eta}_j\right) - \Phi\left(\gamma_{Y_{ij}-1} - \mathbf{B}_i^\top \boldsymbol{\eta}_j\right) \right].$$

After introducing the augmented latent variables $\mathbf{Z}$, the corresponding complete-data measurement likelihood becomes

$$p(\mathbf{Y}, \mathbf{Z} \mid \mathbf{B}, \boldsymbol{\eta}, \boldsymbol{\gamma}) = \prod_{j=1}^{n} \prod_{i=1}^{p} \phi\left(Z_{ij} - \mathbf{B}_i^\top \boldsymbol{\eta}_j\right) I\left\{\gamma_{Y_{ij}-1} < Z_{ij} \le \gamma_{Y_{ij}}\right\},$$

where $\phi(\cdot)$ is the standard normal density. This augmented representation is useful computationally because the factor model becomes Gaussian conditional on $\mathbf{Z}$.

#### S1.2 Positive lower-triangular constraint

The decomposition in Equation (2) is invariant to a rotation of the latent coordinates, since $\boldsymbol{B}\boldsymbol{\eta}_j = (\boldsymbol{B}\boldsymbol{Q})(\boldsymbol{Q}^\top \boldsymbol{\eta}_j)$ for any orthogonal matrix $\boldsymbol{Q}$. To reduce rotational and sign indeterminacy and obtain a stable orientation for posterior computation, we impose a positive lower triangular (PLT) identification constraint on $\boldsymbol{B}$ (Geweke & Zhou, 1996). Specifically,

- the leading $q \times q$ block of $\boldsymbol{B}$ is lower triangular, i.e. $B_{i\ell} = 0$ for $\ell > i$, $i, \ell = 1, \ldots, q$;
- its diagonal entries are strictly positive, $B_{ii} > 0$, $i = 1, \ldots, q$;
- the remaining rows $i = q + 1, \ldots, p$ are unconstrained.

We write $\mathcal{B}_{\mathrm{PLT}}$ for the constrained loading space.

#### S1.3 Nonparametric clustering of latent factors

Cells are clustered through their factor scores $\{\boldsymbol{\eta}_j\}_{j=1}^n$, allowing groups of cells to share a common location–dispersion structure in the latent embedding. A central difficulty is that the number of cell populations is unknown and should be inferred from the data. A Bayesian framework provides a natural mechanism for jointly estimating the number of clusters and the cluster memberships.

The Chinese Restaurant Process (CRP) is a popular prior over the clustering configuration

$$\boldsymbol{\xi} = (\xi_1, \ldots, \xi_n)$$

that accommodates an unknown number of clusters. Under the CRP, the cluster label for cell $j$ follows

$$\Pr(\xi_j = k \mid \xi_1, \ldots, \xi_{j-1}) \propto \begin{cases} |k|, & \text{assigning to an existing cluster labeled } k, \\ \alpha, & \text{assigning to a new cluster,} \end{cases}$$

where $|k|$ is the current size of cluster $k$ and $\alpha > 0$ is the concentration parameter. Although flexible, CRP-based mixture models may generate small extraneous clusters, leading to inconsistent estimation of the number of mixture components even as n goes to infinity. (Miller & Harrison, 2018).

To address this issue, we adopt the mixture of finite mixtures (MFM) framework, which places an explicit prior on the number of components and thereby discourages the creation of extraneous clusters (Miller & Harrison, 2018). The generative process for the factor scores is

$$K \sim p_K(\cdot), \qquad K \in \{1, 2, 3, \ldots\},$$

$$\boldsymbol{\pi} \mid K \sim \text{Dirichlet}(\alpha, \ldots, \alpha),$$

$$\xi_j \mid \boldsymbol{\pi} \sim \text{Categorical}(\boldsymbol{\pi}), \qquad j = 1, \ldots, n,$$

$$(\boldsymbol{\mu}_k, \boldsymbol{\Sigma}_k) \sim \text{NIW}(\boldsymbol{m}_0, \kappa_0, \nu_0, \boldsymbol{S}_0), \qquad k = 1, \ldots, K,$$

and

$$\boldsymbol{\eta}_j \mid \xi_j = k, \boldsymbol{\mu}_k, \boldsymbol{\Sigma}_k \sim N_q(\boldsymbol{\mu}_k, \boldsymbol{\Sigma}_k).$$

Here $p_K(\cdot)$ is a proper probability mass function on the positive integers, and $\text{NIW}(\boldsymbol{m}_0, \kappa_0, \nu_0, \boldsymbol{S}_0)$ denotes the Normal–Inverse-Wishart prior. Specifically,

$$\boldsymbol{\Sigma}_k \sim \text{Inv-Wishart}(\nu_0, \boldsymbol{S}_0),$$

and

$$\boldsymbol{\mu}_k \mid \boldsymbol{\Sigma}_k \sim N_q\left(\boldsymbol{m}_0, \kappa_0^{-1}\boldsymbol{\Sigma}_k\right).$$

Each cluster therefore has its own location $\boldsymbol{\mu}_k$ and dispersion $\boldsymbol{\Sigma}_k$. In particular, the covariance matrix is cluster-specific, allowing different cell populations to vary both in their locations within the latent factor space and in their within-cluster dispersion and dependence structures.

After integrating out the mixture weights $\boldsymbol{\pi}$, the MFM induces an exchangeable partition probability function (Miller & Harrison, 2018). Let

$$n_{k,-j} = \sum_{\ell \neq j} \mathbf{1}(\xi_\ell = k)$$

denote the number of cells assigned to cluster $k$ after excluding cell $j$, and let $K_{+,-j}$ denote the number of occupied clusters in

$$\boldsymbol{\xi}_{-j} = \{\xi_\ell : \ell \neq j\}.$$

The conditional allocation probability follows a Pólya-urn representation. The MFM prior contribution is

$$\Pr(\xi_j = k \mid \boldsymbol{\xi}_{-j}) \propto \begin{cases} n_{k,-j} + \alpha, & \text{for an existing cluster } k, \\ \alpha \dfrac{V_n(K_{+,-j}+1)}{V_n(K_{+,-j})}, & \text{for a new cluster.} \end{cases}$$

The coefficients $V_n(t)$ are defined by

$$V_n(t) = \sum_{K=t}^{\infty} \frac{K_{(t)}}{(\alpha K)^{(n)}} p_K(K),$$

where

$$K_{(t)} = K(K-1)\cdots(K-t+1) = \frac{K!}{(K-t)!}$$

is the falling factorial, and

$$(\alpha K)^{(n)} = (\alpha K)(\alpha K+1)\cdots(\alpha K+n-1) = \frac{\Gamma(\alpha K+n)}{\Gamma(\alpha K)}$$

is the rising factorial, with the convention $K_{(0)} = 1$.

Relative to the CRP, the factor

$$\frac{V_n(K_{+,-j}+1)}{V_n(K_{+,-j})}$$

modifies the prior probability of opening a new cluster. This MFM-specific adjustment controls the proliferation of small extraneous clusters and provides a direct prior on the finite number of mixture components.

#### S1.4 Complete hierarchical model

Combining the ordinal probit factor model with the MFM prior on latent factor scores, the proposed hierarchical ordinal factor model is

$$\begin{aligned}
Y_{ij} = r \iff \gamma_{r-1} < Z_{ij} \le \gamma_r, & \qquad i = 1,\dots,p,\ j = 1,\dots,n, \\
\boldsymbol{Z}_j = \boldsymbol{B}\boldsymbol{\eta}_j + \boldsymbol{\varepsilon}_j, \qquad \boldsymbol{\varepsilon}_j \sim N_p(\boldsymbol{0}, \boldsymbol{I}_p), & \quad j = 1,\dots,n, \\
\boldsymbol{\eta}_j \mid \xi_j = k \sim N_q(\boldsymbol{\mu}_k, \boldsymbol{\Sigma}_k), & \\
(\boldsymbol{\mu}_k, \boldsymbol{\Sigma}_k) \sim \mathrm{NIW}(\boldsymbol{m}_0, \kappa_0, \nu_0, \boldsymbol{S}_0), & \qquad k = 1,\dots,K, \\
\{B_{i\ell} : (i,\ell) \in \mathcal{F}_B\} \overset{\text{ind}}{\sim} N(0, \sigma_B^2), \qquad B_{ii} > 0, & \qquad i = 1,\dots,q, \\
p(\gamma_2,\dots,\gamma_{C-1}) \propto \mathbf{1}\{0 < \gamma_2 < \cdots < \gamma_{C-1}\}, &
\end{aligned}$$

with $\gamma_0 = -\infty$, $\gamma_C = +\infty$, $\gamma_1 = 0$ fixed, and unit latent error variance. Here

$$\mathcal{F}_B = \{(i,\ell) : 1 \le \ell \le i \le q\} \cup \{(i,\ell) : i > q,\ 1 \le \ell \le q\},$$

denotes the set of free loading-matrix entries, while $B_{i\ell} = 0$ for $1 \le i < \ell \le q$. The Gaussian priors on the diagonal elements $B_{ii}$, $i \le q$, are restricted to the positive half-line, $\boldsymbol{\xi} = (\xi_1,\dots,\xi_n)$, $K_+$ is the number of distinct values among the $\xi_j$, and $\mathrm{MFM}(\alpha, p_K)$ denotes the partition law induced by Equation (3). In addition, a graphical representation of the complete hierarchical model is shown in Figure 1.

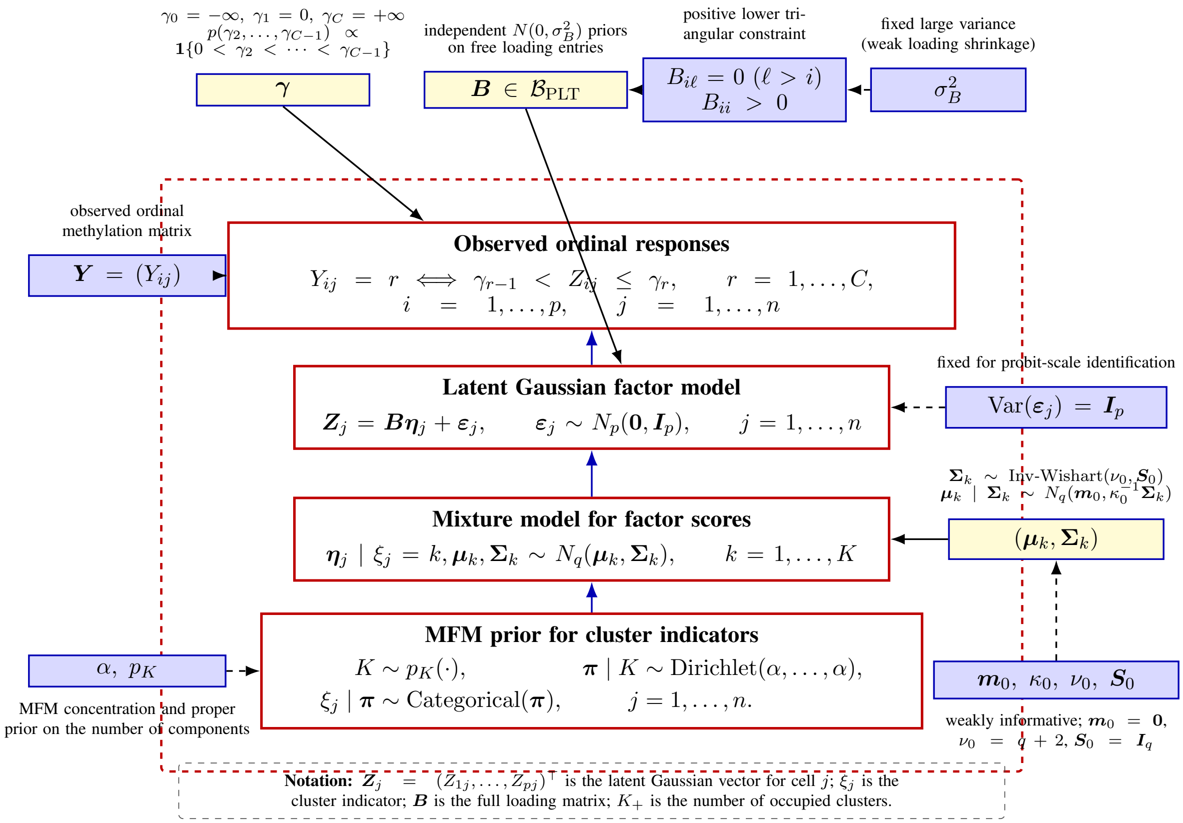

Figure 1: Hierarchical representation of the Bayesian ordinal factor Mixture of Finite Mixtures model. The ordinal observations are linked to latent Gaussian variables through an Albert–Chib threshold model; the latent Gaussian profiles are represented by a low-dimensional factor model; and the factor scores are clustered using an MFM prior with cluster-specific Normal–Inverse-Wishart parameters.

#### S1.5 Initialization and Hyperparameter Specification

We used the following initialization scheme and hyperparameter settings for all simulation studies. Unless otherwise stated, the latent factor dimension was fixed at $q = 10$.

**Initialization** The cutpoints were initialized as

$$\gamma^{(0)} = (-\infty,\ 0,\ 0.5,\ 1.0,\ 1.5,\ \infty),$$

where $\gamma_1 = 0$ was fixed for identifiability and the remaining interior cutpoints were subsequently updated subject to the ordering constraints.

The loading matrix B was initialized under the positive lower triangular identification constraint. For the first $q$ rows, entries below the diagonal were independently generated from

$$B_{ij}^{(0)} \sim N(0, 0.5^2), \qquad j < i,$$

the diagonal entries were initialized as

$$B_{ii}^{(0)} = |X_i|, \qquad X_i \sim N(1, 0.3^2),$$

and entries above the diagonal were set to zero. For rows $i > q$, all $q$ loading coefficients were independently initialized from $N(0, 0.5^2)$.

Given the observed ordinal response $Y_{ij} = c$, the initial latent Gaussian variable was independently generated as

$$Z_{ij}^{(0)} \sim N(0,1)\mathbf{1}\left\{\gamma_{c-1}^{(0)} < Z_{ij}^{(0)} < \gamma_c^{(0)}\right\}.$$

The initial factor scores were then obtained from the regularized least squares projection

$$\boldsymbol{\eta}^{(0)} = \left\{(\mathbf{B}^{(0)})^\top \mathbf{B}^{(0)} + 0.01\mathbf{I}_q\right\}^{-1} (\mathbf{B}^{(0)})^\top \mathbf{Z}^{(0)}.$$

To initialize the mixture allocation, $K_{\text{init}} = 10$ clusters were obtained by applying $K$ means clustering to the columns of $\boldsymbol{\eta}^{(0)}$. For each initial cluster containing at least two observations, the cluster mean was initialized by the empirical mean of its factor scores, and the cluster covariance matrix was initialized by

$$\boldsymbol{\Sigma}_k^{(0)} = \widehat{\text{Cov}}\left(\boldsymbol{\eta}_j^{(0)} : \xi_j^{(0)} = k\right) + 10^{-4}\mathbf{I}_q.$$

For a singleton cluster, its initial mean was set equal to the corresponding factor score, with a regularized covariance matrix used to ensure positive definiteness. These quantities were used only to initialize the Markov chain and did not determine the prior specification described below.

**Prior hyperparameters** Under the prior specification described in Section S1.4, we set $\sigma_B = 5$, $\mathbf{m}_0 = \mathbf{0}_q$, $\kappa_0 = 0.1$, $\nu_0 = q + 2$, and $\mathbf{S}_0 = \mathbf{I}_q$. Under the inverse-Wishart parameterization used in the sampler, these settings imply $E(\boldsymbol{\Sigma}_k) = \mathbf{I}_q$. For the MFM prior, we used $K - 1 \sim \text{Poisson}(4)$

and set the symmetric Dirichlet concentration parameter to $\alpha = 1$. The summation used to evaluate $V_n(t)$ was truncated at $K = 200$, with at most 40 occupied clusters maintained during sampling.

**MCMC settings and convergence assessment** For each simulation replicate, one MCMC chain was run for 2,000 iterations, with the first 1,000 iterations discarded as burn-in. For each real-data analysis, one chain was run for 5,000 iterations, with the first 2,500 iterations discarded as burn-in. In the simulation studies, convergence of the clustering configuration was assessed using post-burn-in trace plots of the adjusted Rand index (ARI). No additional thinning was applied.

#### S1.6 Posterior Clustering Summary

Because the partition $\boldsymbol{\xi}$ is a combinatorial object, it is not naturally summarized by a posterior mean or mode. We summarize the posterior clustering using the least-squares partition estimator based on the posterior membership matrices (Dahl, 2006). For each post-burn-in iteration $t = 1, \ldots, M$, define the $n \times n$ membership matrix

$$\boldsymbol{\Delta}^{(t)} = \big[\mathbf{1}\{\xi_i^{(t)} = \xi_j^{(t)}\}\big]_{i,j\in\{1,\ldots,n\}},$$

where $\boldsymbol{\Delta}^{(t)}(i,j) = 1$ indicates that cells $i$ and $j$ share a cluster in the $t$-th posterior sample. The element-wise average

$$\overline{\boldsymbol{\Delta}} = \frac{1}{M}\sum_{t=1}^{M}\boldsymbol{\Delta}^{(t)}$$

summarizes the posterior co-clustering probabilities among cells, with $\overline{\boldsymbol{\Delta}}(i,j)$ estimating the posterior probability that cells $i$ and $j$ belong to the same cluster. The Dahl estimate is the sampled partition closest to $\overline{\boldsymbol{\Delta}}$ in squared distance,

$$C_{\mathrm{LS}} = \underset{t\in\{1,\ldots,M\}}{\arg\min}\sum_{i=1}^{n}\sum_{j=1}^{n}\big\{\boldsymbol{\Delta}^{(t)}(i,j) - \overline{\boldsymbol{\Delta}}(i,j)\big\}^2.$$

The representative partition corresponding to $C_{\mathrm{LS}}$ is used for the clustering results reported in the simulation and real-data analyses. This estimator incorporates information from all retained posterior partitions through the posterior co-clustering matrix while selecting a valid partition that actually occurred in the MCMC sample.

#### S2 Full conditional distributions and their derivations

**Latent Gaussian variables** From the latent Gaussian factor model,

$$Z_{ij} \mid \boldsymbol{B}, \boldsymbol{\eta}_j \sim N(\boldsymbol{B}_{i\cdot}\boldsymbol{\eta}_j, 1)\,.$$

If $Y_{ij} = r$, ordinal consistency additionally requires $\gamma_{r-1} < Z_{ij} \leq \gamma_r$. Hence the conditional kernel is

$$p(Z_{ij} \mid \text{rest}) \propto \phi(Z_{ij} - \boldsymbol{B}_{i\cdot}\boldsymbol{\eta}_j)\,\mathbf{1}\left\{\gamma_{r-1} < Z_{ij} \leq \gamma_r\right\}.$$

Therefore,

$$Z_{ij} \mid \text{rest} \sim \text{TN}\big(\boldsymbol{B}_{i\cdot}\boldsymbol{\eta}_j, 1; \gamma_{Y_{ij}-1}, \gamma_{Y_{ij}}\big)\,.$$

**Factor scores** Suppose cell $j$ is currently assigned to cluster $k = \xi_j$. The measurement model and mixture prior are

$$\boldsymbol{Z}_j \mid \boldsymbol{\eta}_j, \boldsymbol{B} \sim N_p(\boldsymbol{B}\boldsymbol{\eta}_j, \boldsymbol{I}_p), \qquad \boldsymbol{\eta}_j \mid \xi_j = k, \boldsymbol{\mu}_k, \boldsymbol{\Sigma}_k \sim N_q(\boldsymbol{\mu}_k, \boldsymbol{\Sigma}_k).$$

Thus,

$$p(\boldsymbol{\eta}_j \mid \text{rest}) \propto \exp\left\{-\frac{1}{2}(\boldsymbol{Z}_j - \boldsymbol{B}\boldsymbol{\eta}_j)^\top(\boldsymbol{Z}_j - \boldsymbol{B}\boldsymbol{\eta}_j)\right\}$$
$$\times \exp\left\{-\frac{1}{2}(\boldsymbol{\eta}_j - \boldsymbol{\mu}_k)^\top\boldsymbol{\Sigma}_k^{-1}(\boldsymbol{\eta}_j - \boldsymbol{\mu}_k)\right\}.$$

Keeping only terms involving $\boldsymbol{\eta}_j$, the conditional kernel becomes

$$p(\boldsymbol{\eta}_j \mid \text{rest}) \propto \exp\left\{-\frac{1}{2}\left[\boldsymbol{\eta}_j^\top(\boldsymbol{B}^\top\boldsymbol{B} + \boldsymbol{\Sigma}_k^{-1})\boldsymbol{\eta}_j - 2\boldsymbol{\eta}_j^\top(\boldsymbol{B}^\top\boldsymbol{Z}_j + \boldsymbol{\Sigma}_k^{-1}\boldsymbol{\mu}_k)\right]\right\}.$$

Completing the square gives

$$\boldsymbol{\eta}_j \mid \text{rest} \sim N_q(\boldsymbol{m}_j^*, \boldsymbol{V}_j^*),$$

where

$$\boldsymbol{V}_j^* = (\boldsymbol{B}^\top \boldsymbol{B} + \boldsymbol{\Sigma}_k^{-1})^{-1}, \qquad \boldsymbol{m}_j^* = \boldsymbol{V}_j^* (\boldsymbol{B}^\top \boldsymbol{Z}_j + \boldsymbol{\Sigma}_k^{-1} \boldsymbol{\mu}_k).$$

**Factor loadings with unconstrained rows** For $i > q$, the likelihood contribution for row $i$ is

$$Z_{ij} = \boldsymbol{B}_{i\cdot} \boldsymbol{\eta}_j + \epsilon_{ij}, \qquad \epsilon_{ij} \sim N(0, 1),$$

and the free loading elements have independent $N(0, \sigma_B^2)$ priors. Therefore,

$$p(\boldsymbol{B}_{i\cdot}^\top \mid \text{rest}) \propto \exp\left\{ -\frac{1}{2} \sum_{j=1}^n (Z_{ij} - \boldsymbol{B}_{i\cdot} \boldsymbol{\eta}_j)^2 - \frac{1}{2\sigma_B^2} \boldsymbol{B}_{i\cdot} \boldsymbol{B}_{i\cdot}^\top \right\}.$$

Collecting the quadratic and linear terms in $\boldsymbol{B}_{i\cdot}^\top$ gives

$$p(\boldsymbol{B}_{i\cdot}^\top \mid \text{rest}) \propto \exp\left\{ -\frac{1}{2} \left[ \boldsymbol{B}_{i\cdot} \left( \sum_{j=1}^n \boldsymbol{\eta}_j \boldsymbol{\eta}_j^\top + \sigma_B^{-2} \boldsymbol{I}_q \right) \boldsymbol{B}_{i\cdot}^\top - 2 \boldsymbol{B}_{i\cdot} \sum_{j=1}^n Z_{ij} \boldsymbol{\eta}_j \right] \right\}.$$

Hence,

$$\boldsymbol{B}_{i\cdot}^\top \mid \text{rest} \sim N_q(\boldsymbol{m}_i^B, \boldsymbol{V}_i^B),$$

where

$$\boldsymbol{V}_i^B = \left( \sum_{j=1}^n \boldsymbol{\eta}_j \boldsymbol{\eta}_j^\top + \sigma_B^{-2} \boldsymbol{I}_q \right)^{-1}, \qquad \boldsymbol{m}_i^B = \boldsymbol{V}_i^B \sum_{j=1}^n Z_{ij} \boldsymbol{\eta}_j.$$

**Factor loadings with positive lower-triangular rows** For $i \le q$, only the first $i$ entries in row $i$ are free. Let

$$\boldsymbol{H}_j^{(i)} = (\eta_{j1}, \ldots, \eta_{ji})^\top.$$

Applying the same Gaussian calculation to $\boldsymbol{B}_{i,1:i}^\top$ gives the unconstrained kernel

$$p(\boldsymbol{B}_{i,1:i}^\top \mid \text{rest}) \propto \exp\left\{ -\frac{1}{2} \left[ \boldsymbol{B}_{i,1:i} \widetilde{\boldsymbol{V}}_i^{-1} \boldsymbol{B}_{i,1:i}^\top - 2 \boldsymbol{B}_{i,1:i} \widetilde{\boldsymbol{V}}_i^{-1} \widetilde{\boldsymbol{m}}_i \right] \right\},$$

where

$$\widetilde{\boldsymbol{V}}_i = \left( \sum_{j=1}^n \boldsymbol{H}_j^{(i)} \boldsymbol{H}_j^{(i)\top} + \sigma_B^{-2} \boldsymbol{I}_i \right)^{-1},$$

and

$$\widetilde{\boldsymbol{m}}_i = \widetilde{\boldsymbol{V}}_i \sum_{j=1}^{n} Z_{ij} \boldsymbol{H}_j^{(i)}.$$

The PLT identification constraint additionally requires $B_{ii} > 0$. Therefore,

$$\boldsymbol{B}_{i,1:i}^{\top} \mid \text{rest} \sim N_i(\widetilde{\boldsymbol{m}}_i, \widetilde{\boldsymbol{V}}_i) \mathbf{1}\{B_{ii} > 0\}, \qquad i \leq q,$$

while $B_{i\ell} = 0$ for $\ell > i$.

**Cutpoints** For $r = 2, \ldots, C-1$, the cutpoint $\gamma_r$ enters the augmented likelihood only through the ordinal constraints. For observations with $Y_{ij} = r$,

$$Z_{ij} \leq \gamma_r,$$

whereas observations with $Y_{ij} = r+1$ require

$$Z_{ij} > \gamma_r.$$

Together with the ordering constraint $\gamma_{r-1} < \gamma_r < \gamma_{r+1}$, define

$$L_r = \max\left\{\gamma_{r-1}, \max_{Y_{ij}=r} Z_{ij}\right\}, \qquad U_r = \min\left\{\gamma_{r+1}, \min_{Y_{ij}=r+1} Z_{ij}\right\}.$$

Under the flat prior subject to the ordering constraint, the conditional kernel is therefore

$$p(\gamma_r \mid \text{rest}) \propto \mathbf{1}\{L_r < \gamma_r < U_r\}.$$

Hence,

$$\gamma_r \mid \text{rest} \sim \text{Uniform}(L_r, U_r), \qquad r = 2, \ldots, C-1.$$

The cutpoint $\gamma_1 = 0$ remains fixed for identification.

**Cluster allocations** Before updating $\xi_j$, cell $j$ is removed from its current cluster. Let $n_{k,-j}$ denote the size of occupied cluster $k$ after removal, and let $K_{+,-j}$ denote the resulting number of occupied clusters.

For an existing cluster $k \in \mathcal{C}_{-j}$, the MFM partition prior contributes the factor $n_{k,-j} + \alpha$, while the conditional density of the factor score contributes

$$N_q(\boldsymbol{\eta}_j; \boldsymbol{\mu}_k, \boldsymbol{\Sigma}_k).$$

Thus, the allocation kernel is

$$\Pr(\xi_j = k \mid \text{rest}) \propto (n_{k,-j} + \alpha) N_q(\boldsymbol{\eta}_j; \boldsymbol{\mu}_k, \boldsymbol{\Sigma}_k), \qquad k \in \mathcal{C}_{-j}.$$

For a new cluster, the MFM partition prior contributes

$$\alpha \frac{V_n(K_{+,-j} + 1)}{V_n(K_{+,-j})}.$$

Integrating the new component parameters $(\boldsymbol{\mu}, \boldsymbol{\Sigma})$ with respect to the Normal–Inverse-Wishart base distribution gives

$$m_0(\boldsymbol{\eta}_j) = t_{\nu_0 - q + 1}\left(\boldsymbol{\eta}_j; \boldsymbol{m}_0, \frac{\kappa_0 + 1}{\kappa_0(\nu_0 - q + 1)} \boldsymbol{S}_0\right).$$

Therefore,

$$\Pr(\xi_j = \text{new} \mid \text{rest}) \propto \alpha \frac{V_n(K_{+,-j} + 1)}{V_n(K_{+,-j})} m_0(\boldsymbol{\eta}_j).$$

The existing-cluster and new-cluster weights are normalized before sampling the updated allocation.

**Gaussian mixture parameters** Consider an occupied cluster $k$ with $n_k$ factor scores. Define

$$\overline{\boldsymbol{\eta}}_k = \frac{1}{n_k} \sum_{j \in \mathcal{C}_k} \boldsymbol{\eta}_j, \qquad \boldsymbol{S}_k = \sum_{j \in \mathcal{C}_k} (\boldsymbol{\eta}_j - \overline{\boldsymbol{\eta}}_k)(\boldsymbol{\eta}_j - \overline{\boldsymbol{\eta}}_k)^\top.$$

The Gaussian likelihood satisfies

$$\sum_{j \in \mathcal{C}_k} (\boldsymbol{\eta}_j - \boldsymbol{\mu}_k)(\boldsymbol{\eta}_j - \boldsymbol{\mu}_k)^\top = \boldsymbol{S}_k + n_k(\overline{\boldsymbol{\eta}}_k - \boldsymbol{\mu}_k)(\overline{\boldsymbol{\eta}}_k - \boldsymbol{\mu}_k)^\top.$$

Combining the second term with the Normal prior for $\boldsymbol{\mu}_k$ gives

$$n_k(\overline{\boldsymbol{\eta}}_k - \boldsymbol{\mu}_k)(\overline{\boldsymbol{\eta}}_k - \boldsymbol{\mu}_k)^\top + \kappa_0(\boldsymbol{\mu}_k - \boldsymbol{m}_0)(\boldsymbol{\mu}_k - \boldsymbol{m}_0)^\top$$

$$= (\kappa_0 + n_k)(\boldsymbol{\mu}_k - \boldsymbol{m}_k^*)(\boldsymbol{\mu}_k - \boldsymbol{m}_k^*)^\top + \boldsymbol{R}_k,$$

where

$$\boldsymbol{m}_k^* = \frac{\kappa_0 \boldsymbol{m}_0 + n_k \overline{\boldsymbol{\eta}}_k}{\kappa_0 + n_k},$$

and

$$\boldsymbol{R}_k = \frac{\kappa_0 n_k}{\kappa_0 + n_k}(\overline{\boldsymbol{\eta}}_k - \boldsymbol{m}_0)(\overline{\boldsymbol{\eta}}_k - \boldsymbol{m}_0)^\top.$$

Combining these terms with the inverse-Wishart prior gives the Normal–Inverse-Wishart posterior,

$$\boldsymbol{\Sigma}_k \mid \text{rest} \sim \text{IW}\left(\nu_0 + n_k,\ \boldsymbol{S}_0 + \boldsymbol{S}_k + \boldsymbol{R}_k\right),$$

followed by

$$\boldsymbol{\mu}_k \mid \boldsymbol{\Sigma}_k, \text{rest} \sim N_q\left(\frac{\kappa_0 \boldsymbol{m}_0 + n_k \overline{\boldsymbol{\eta}}_k}{\kappa_0 + n_k}, \frac{\boldsymbol{\Sigma}_k}{\kappa_0 + n_k}\right).$$

### S3 Algorithm

We present the Gibbs sampler for ordinal factor MFM in Algorithm 1.

### S4 Tuning Parameters and Benchmark Settings

#### S4.1 Gap statistics

For benchmark clustering methods that require the number of clusters to be specified in advance, including K-means, hierarchical clustering, and Gaussian mixture models, we selected the cluster number using the gap statistic. For a candidate number of clusters $K$, the gap statistic is defined as

$$\text{Gap}(K) = \mathbb{E}^*\{\log W_K\} - \log W_K,$$

**Algorithm 1:** Gibbs sampler for the ordinal factor MFM model

**procedure** Ordinal-MFM

Initialize $\boldsymbol{Z}, \boldsymbol{B}, \{\boldsymbol{\eta}_j\}_{j=1}^n, \boldsymbol{\gamma}, \boldsymbol{\xi}$, and $\{(\boldsymbol{\mu}_k, \boldsymbol{\Sigma}_k)\}_{k=1}^{K_+}$.

**for** each iter = 1 to M **do**

Update $Z_{ij}$, $i = 1, \ldots, p$, $j = 1, \ldots, n$:

$$Z_{ij} \mid \text{rest} \sim \text{TN}\big(\boldsymbol{B}_{i\cdot}\boldsymbol{\eta}_j, 1; \gamma_{Y_{ij}-1}, \gamma_{Y_{ij}}\big)\,.$$

Update $\boldsymbol{\eta}_j$, $j = 1, \ldots, n$, with $k = \xi_j$:

$$\boldsymbol{\eta}_j \mid \text{rest} \sim N_q(\boldsymbol{m}_j^*, \boldsymbol{V}_j^*), \quad \boldsymbol{V}_j^* = (\boldsymbol{B}^\top \boldsymbol{B} + \boldsymbol{\Sigma}_k^{-1})^{-1}, \quad \boldsymbol{m}_j^* = \boldsymbol{V}_j^*(\boldsymbol{B}^\top \boldsymbol{Z}_j + \boldsymbol{\Sigma}_k^{-1}\boldsymbol{\mu}_k).$$

Update $\boldsymbol{B}$ row by row. For $i > q$,

$$\boldsymbol{B}_{i\cdot}^\top \mid \text{rest} \sim N_q(\boldsymbol{m}_i^B, \boldsymbol{V}_i^B), \quad \boldsymbol{V}_i^B = \left(\sum_{j=1}^n \boldsymbol{\eta}_j \boldsymbol{\eta}_j^\top + \sigma_B^{-2}\boldsymbol{I}_q\right)^{-1}, \quad \boldsymbol{m}_i^B = \boldsymbol{V}_i^B \sum_{j=1}^n Z_{ij}\boldsymbol{\eta}_j.$$

For $i \le q$, letting $\boldsymbol{H}_j^{(i)} = (\eta_{j1}, \ldots, \eta_{ji})^\top$,

$$\boldsymbol{B}_{i,1:i}^\top \mid \text{rest} \sim N_i(\widetilde{\boldsymbol{m}}_i, \widetilde{\boldsymbol{V}}_i)\mathbf{1}\{B_{ii} > 0\}, \quad \widetilde{\boldsymbol{V}}_i = \left(\sum_{j=1}^n \boldsymbol{H}_j^{(i)}\boldsymbol{H}_j^{(i)\top} + \sigma_B^{-2}\boldsymbol{I}_i\right)^{-1}, \quad \widetilde{\boldsymbol{m}}_i = \widetilde{\boldsymbol{V}}_i \sum_{j=1}^n Z_{ij}\boldsymbol{H}_j^{(i)}.$$

Update $\gamma_r$, $r = 2, \ldots, C-1$:

$$\gamma_r \mid \text{rest} \sim \text{Uniform}(L_r, U_r), \quad L_r = \max\left\{\gamma_{r-1}, \max_{Y_{ij}=r} Z_{ij}\right\}, \quad U_r = \min\left\{\gamma_{r+1}, \min_{Y_{ij}=r+1} Z_{ij}\right\}.$$

Update $\xi_j$, $j = 1, \ldots, n$, after removing $j$ from its current cluster:

$$\Pr(\xi_j = k \mid \text{rest}) \propto \begin{cases} (n_{k,-j} + \alpha) N_q(\boldsymbol{\eta}_j; \boldsymbol{\mu}_k, \boldsymbol{\Sigma}_k), & k \in \mathcal{C}_{-j}, \\ \alpha \dfrac{V_n(K_{+,-j}+1)}{V_n(K_{+,-j})} t_{\nu_0 - q + 1}\left(\boldsymbol{\eta}_j; \boldsymbol{m}_0, \dfrac{\kappa_0 + 1}{\kappa_0(\nu_0 - q + 1)}\boldsymbol{S}_0\right), & k \text{ is new.} \end{cases}$$

Update $(\boldsymbol{\mu}_k, \boldsymbol{\Sigma}_k)$ for each occupied cluster $k$:

$$\boldsymbol{\Sigma}_k \mid \text{rest} \sim \text{IW}\left(\nu_0 + n_k,\ \boldsymbol{S}_0 + \boldsymbol{S}_k + \boldsymbol{R}_k\right), \qquad \boldsymbol{\mu}_k \mid \boldsymbol{\Sigma}_k, \text{rest} \sim N_q\left(\frac{\kappa_0 \boldsymbol{m}_0 + n_k \overline{\boldsymbol{\eta}}_k}{\kappa_0 + n_k}, \frac{\boldsymbol{\Sigma}_k}{\kappa_0 + n_k}\right),$$

where

$$\overline{\boldsymbol{\eta}}_k = n_k^{-1} \sum_{j \in \mathcal{C}_k} \boldsymbol{\eta}_j, \qquad \boldsymbol{S}_k = \sum_{j \in \mathcal{C}_k} (\boldsymbol{\eta}_j - \overline{\boldsymbol{\eta}}_k)(\boldsymbol{\eta}_j - \overline{\boldsymbol{\eta}}_k)^\top,$$

and

$$\boldsymbol{R}_k = \frac{\kappa_0 n_k}{\kappa_0 + n_k}(\overline{\boldsymbol{\eta}}_k - \boldsymbol{m}_0)(\overline{\boldsymbol{\eta}}_k - \boldsymbol{m}_0)^\top.$$

**end for**

**end procedure**

where $W_K$ denotes the within-cluster dispersion for $K$ clusters and $\mathbb{E}^*\{\log W_K\}$ is the expectation under a reference null distribution generated by resampling from the data domain. This criterion selects the value of $K$ whose observed clustering structure deviates most strongly from that expected under the reference distribution.

#### S4.2 Hyper-parameter

For methods involving dimension reduction, the reduced dimension was fixed at ten throughout the benchmark analyses. In the proposed *SMORE* model, the corresponding latent dimension is denoted by $q$; accordingly, we set $q = 10$ when constructing its low-dimensional latent representation.This choice ensured a fair comparison by giving all methods a common representation size and avoiding additional method-specific tuning. Larger latent dimensions may provide additional flexibility, but they also introduce more parameters and can make posterior computation more demanding. Because runtime was not the primary focus of this study, we fixed the dimension at 10 as a practical compromise rather than treating it as an optimized tuning parameter. We additionally conducted a sensitivity analysis using $q \in \{5, 10, 15, 20\}$ to evaluate the robustness of *SMORE* to the choice of latent dimension; the results are reported in Supplementary Section S8.5.

### S5 Evaluation metrics

To evaluate clustering performance on the real single-cell methylation datasets, we introduce the following standard external validation metrics:

**Adjusted Rand Index (ARI)** measures the agreement between the predicted clustering and the reference cell-type labels based on pairwise assignments. Unlike the Rand index, ARI corrects for the agreement expected by chance. It is defined as

$$\text{ARI} = \frac{\sum_{ij}\binom{n_{ij}}{2} - \left[\sum_i \binom{a_i}{2} \sum_j \binom{b_j}{2}\right] / \binom{n}{2}}{\frac{1}{2}\left[\sum_i \binom{a_i}{2} + \sum_j \binom{b_j}{2}\right] - \left[\sum_i \binom{a_i}{2} \sum_j \binom{b_j}{2}\right] / \binom{n}{2}},$$

where $n_{ij}$ is the number of cells shared by true class $i$ and predicted cluster $j$, $a_i = \sum_j n_{ij}$, $b_j = \sum_i n_{ij}$, and $n$ is the total number of cells. Larger ARI values indicate stronger agreement with the reference labels.

**Normalized Mutual Information (NMI)** quantifies the shared information between the predicted cluster labels and the true cell-type labels, normalized to account for the entropy of the two labelings:

$$\text{NMI}(U, V) = \frac{I(U; V)}{\sqrt{H(U)H(V)}},$$

where $U$ denotes the true labels, $V$ denotes the predicted clusters, $I(U;V)$ is their mutual information, and $H(\cdot)$ is entropy. NMI ranges from 0 to 1, with higher values indicating better correspondence between cluster assignments and cell types.

**Adjusted Mutual Information (AMI)** is an adjusted version of mutual information that accounts for chance agreement between two labelings:

$$\mathrm{AMI}(U,V) = \frac{I(U;V) - \mathbb{E}\{I(U;V)\}}{\mathrm{avg}\{H(U),H(V)\} - \mathbb{E}\{I(U;V)\}}.$$

AMI is particularly useful when comparing clusterings with different numbers of clusters, because it penalizes agreement that may arise solely from random partitions.

**Homogeneity** evaluates whether each predicted cluster contains cells from only a single reference cell type:

$$\text{Homogeneity} = 1 - \frac{H(U \mid V)}{H(U)}.$$

A value close to 1 indicates that predicted clusters are internally pure with respect to the true cell-type labels.

**Completeness** evaluates whether cells from the same reference cell type are assigned to the same predicted cluster:

$$\text{Completeness} = 1 - \frac{H(V \mid U)}{H(V)}.$$

A value close to 1 indicates that each true cell type is well recovered without being split across multiple predicted clusters.

### S6 Model-based simulation design

**Simulation settings** The primary model-based simulation was fully synthetic and generated independently of the real methylation datasets. We fixed the true number of clusters at $K_0 = 5$, the number of genomic features at $p = 5000$, and the generative factor dimension at $q_0 = 4$. Sample sizes were $n \in \{500, 1000, 2000, 4000\}$. For each sample size, balanced and unbalanced cluster configurations were crossed with strong and weak signal regimes. Balanced clusters had equal sizes, whereas the unbalanced design used target proportions $(0.40, 0.30, 0.15, 0.10, 0.05)$. Twenty replicate datasets were generated for each setting.

**Latent cluster structure** Let $g_j \in \{1, \ldots, K_0\}$ denote the true cluster label of cell $j$ and let $\pi_k = n_k/n$ denote the corresponding cluster proportion. Cluster centers were constructed from Helmert contrasts. If $\mathbf{h}_k^\top$ is the $k$th row of the $K_0 \times (K_0 - 1)$ Helmert contrast matrix, define

$$\mathbf{c}_k = \frac{\mathbf{h}_k}{\|\mathbf{h}_k\|_2}, \qquad \overline{\mathbf{c}} = \sum_{k=1}^{K_0} \pi_k \mathbf{c}_k, \qquad \boldsymbol{\nu}_k = \delta(\mathbf{c}_k - \overline{\mathbf{c}}),$$

with $\delta = 1$. This construction gives distinct, centered cluster directions without imposing spherical within-cluster geometry.

For each cluster, an independent random orthogonal matrix $\mathbf{Q}_k$ was obtained from the QR decomposition of a $q_0 \times q_0$ matrix with independent standard normal entries. For anisotropy parameter $a > 1$, define

$$\mathbf{d}(a) = \left(a,\, a^{-1/(q_0-1)}, \ldots, a^{-1/(q_0-1)}\right)^\top, \qquad \widetilde{\mathbf{d}}(a) = \frac{\mathbf{d}(a)}{\left\{\prod_{\ell=1}^{q_0} d_\ell(a)\right\}^{1/q_0}}.$$

The cluster-specific covariance matrix was

$$\boldsymbol{\Sigma}_k = \sigma_\eta^2 v_k \, \mathbf{Q}_k \, \mathrm{diag}\{\widetilde{\mathbf{d}}(a)\} \mathbf{Q}_k^\top,$$

where $(v_1, \ldots, v_5) = (0.6, 0.8, 1.0, 1.4, 2.0)$ are cluster-specific covariance-scale multipliers. Latent factor scores were then sampled as

$$\boldsymbol{\eta}_j^0 \mid g_j = k \sim N_{q_0}(\boldsymbol{\nu}_k, \boldsymbol{\Sigma}_k).$$

The random orientations $\mathbf{Q}_k$ were regenerated within each replicate.

**Feature-level signal and ordinal observations** For each feature, an initial baseline was drawn independently as

$$\alpha_i \sim N(0.85, 0.20^2).$$

A subset $\mathcal{S}$ of $\lfloor p_{\text{sig}} p \rceil$ informative features was sampled uniformly without replacement. For $i \in \mathcal{S}$, the loading vector was generated as $\boldsymbol{\lambda}_i \sim N_{q_0}(\mathbf{0}, \mathbf{I}_{q_0})$ and the baseline was set to $\alpha_i = 0$; for $i \notin \mathcal{S}$, $\boldsymbol{\lambda}_i = \mathbf{0}$. Conditional on the latent factor scores, the latent Gaussian measurements followed

$$\widetilde{Z}_{ij} = \alpha_i + s\, \boldsymbol{\lambda}_i^\top \boldsymbol{\eta}_j^0 + \epsilon_{ij}, \qquad \epsilon_{ij} \overset{\text{iid}}{\sim} N(0, 1).$$

The strong and weak regimes used the following parameters:

| Regime | $s$ | $p_{\text{sig}}$ | $\sigma_\eta$ | $a$ |
|---|---|---|---|---|
| Strong | 5.0 | 0.40 | 0.20 | 8 |
| Weak | 4.0 | 0.25 | 0.24 | 10 |

The center scaling parameter was fixed at $\delta = 1$ in both regimes. Thus, the two regimes differed through the observation-level signal, the fraction of informative features, within-cluster dispersion and covariance anisotropy rather than through a change in the cluster-center locations.

To obtain five ordinal categories, four replicate-specific cutpoints were defined as the empirical 0.15, 0.25, 0.40 and 0.60 quantiles of all entries of $\widetilde{\mathbf{Z}}$. Each $\widetilde{Z}_{ij}$ was then assigned to its corresponding interval, producing an ordinal matrix $\mathbf{Y} \in \{1, \ldots, 5\}^{p \times n}$ with approximate marginal category proportions

$$(0.15, 0.10, 0.15, 0.20, 0.40).$$

After generation, cell columns and the corresponding true labels were permuted using the same setting-specific random permutation. True labels were used only for external performance evaluation.

### S7 Distribution-mixture simulation

**Feature-level mixture model** GSE152011 was used to estimate the feature-level distributions employed in the simulation data-generating mechanism. This dataset was used only for simulation calibration and was not included in the real-data clustering benchmarks. Entries with zero coverage were treated as missing, and methylation proportions were computed as ratios of methylated counts to coverage. Observed values were partitioned into two subsets: (i) edge values near 0 or 1, representing U-shaped behavior, and (ii) mid-range values between 0.2 and 0.8, representing dense variability.

For each subset, Beta distribution parameters were estimated via moment matching. Let $\mu = \mathbb{E}(X)$ and $v = \mathrm{Var}(X)$. The parameters are given by

$$\alpha = \mu \left( \frac{\mu(1-\mu)}{v} - 1 \right), \quad \beta = (1-\mu) \left( \frac{\mu(1-\mu)}{v} - 1 \right).$$

This yields two representative components: a U-shaped distribution capturing extreme values and a dense distribution capturing moderate variability.

Each feature was generated from a mixture of four components: a U-shaped Beta distribution, a dense Beta distribution, a point mass at 1 (one-inflation), and a point mass at 0 (zero-inflation). Formally, each simulated value $x$ follows

$$x \sim w_u \cdot \mathrm{Beta}(\alpha_u, \beta_u) + w_d \cdot \mathrm{Beta}(\alpha_d, \beta_d) + w_1 \cdot \delta_1 + w_0 \cdot \delta_0,$$

where $(w_u, w_d, w_1, w_0)$ are mixture weights satisfying $w_u + w_d + w_1 + w_0 = 1$. This formulation captures both zero/one inflation and heavy-tailed behavior commonly observed in methylation data.

**Simulation regimes** Cluster structure was generated by assigning different mixture weights to latent clusters while keeping the underlying Beta components fixed. Consequently, clusters differ in their overall distributional shapes rather than solely in mean levels, creating a challenging clustering problem that resembles heterogeneous biological data.

We considered four simulation regimes by crossing cluster balance with signal strength: balanced strong, balanced weak, unbalanced strong, and unbalanced weak. In the strong-signal setting, cluster-specific mixture weights were chosen to induce clearer separation among methylation profiles. In the weak-signal setting, mixture weights were more similar across clusters, resulting in greater overlap. Balanced settings used equal cluster sizes, whereas unbalanced settings used unequal cluster sizes. Each dataset contained $K = 5$ clusters and $p = 5000$ CpG features. The main benchmark used sample sizes $n \in \{500, 1000, 2000, 4000\}$, with 20 independent replicates generated for each scenario.

**Data construction** To reflect common preprocessing steps in epigenetic analysis, continuous values in $[0, 1]$ were discretized into ordinal categories. Let $0 = c_0 < c_1 < \cdots < c_L = 1$ denote equally spaced cut points. Each value $x$ was mapped to an ordinal level:

$$x^{(\text{ord})} = \ell \quad \text{if } x \in (c_{\ell-1}, c_\ell].$$

In the experiments, $L = 5$ levels were used. To eliminate ordering artifacts, all samples were randomly permuted after generation. The corresponding cluster labels were shuffled accordingly and used solely for evaluation.

The simulation framework is grounded in real data, incorporates mixture-based heterogeneity, accounts for zero/one inflation, and evaluates both strong and weak signal regimes under varying cluster configurations. This design provides a realistic and challenging benchmark for clustering methods.

#### S7.1 Result

Figure 2 summarizes the clustering performance across the twelve simulation settings, and Table 1 reports the corresponding numerical results.

Overall, *SMORE* recovered the latent clustering structure most reliably in the strong-signal settings. In the unbalanced strong scenarios, *SMORE* achieved consistently high ARI values across sample sizes, with mean ARI equal to 0.918, 0.905, and 0.906 for $n = 500, 1000$, and $2000$, respectively. These results were comparable to or better than the best benchmark pipelines, while avoiding the more variable behavior observed for several dimension-reduction-plus-clustering combinations. In the balanced strong scenarios, *SMORE* also performed well, with mean ARI increasing from 0.758 at $n = 500$ to 0.847 at $n = 1000$ and remaining high at 0.807 for $n = 2000$.

As expected, the weak-signal settings were more challenging for all methods, and especially for the proposed model-based approach. *SMORE* showed reduced ARI in the balanced weak scenarios, with mean ARI ranging from 0.381 to 0.433, indicating that weaker between-cluster separation limits exact recovery of the true partition. This behavior reflects the construction of this simulation setting: the weak-signal data primarily reduce marginal separation between clusters, so the latent groups differ only subtly relative to within-cluster variability. Under this type of stress test,

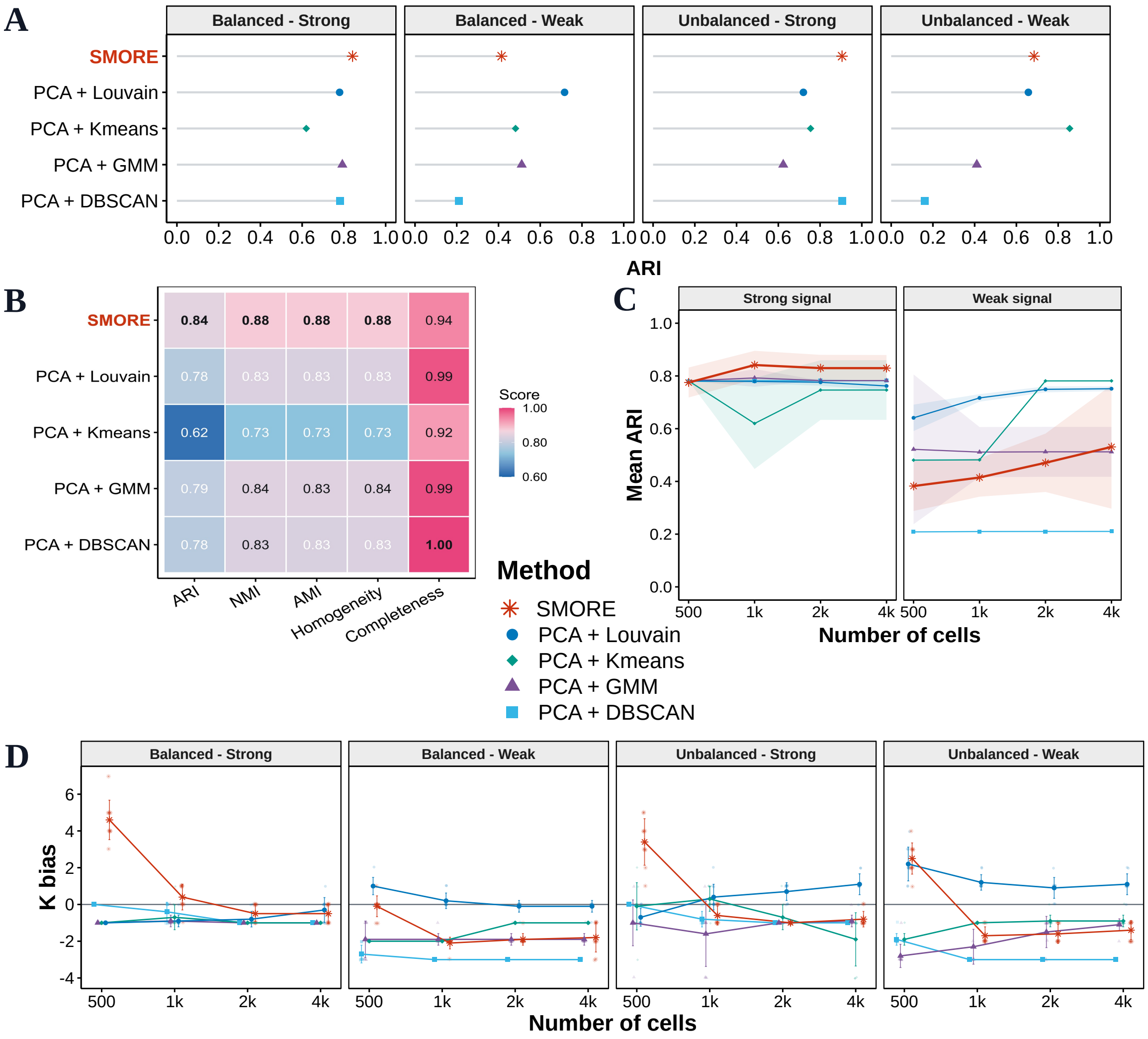

Figure 2: Performance comparison under the distribution-mixture simulations. **(A)** ARI at $n = 1{,}000$ cells across four simulation scenarios defined by balanced or unbalanced cluster sizes and strong or weak signal. **(B)** Performance in terms of ARI, NMI, AMI, homogeneity, and completeness under the balanced strong-signal setting at $n = 1{,}000$ cells. **(C)** Scalability as the sample size increases under the balanced strong- and weak-signal settings. Lines indicate mean ARI, and shaded ribbons indicate $\pm$ one standard deviation across replicates. *SMORE* is highlighted in red.

Table 1: Distribution-mixture simulation performance of *SMORE* and the top 10 benchmark pairs across sample sizes and scenarios. For each scenario and sample size, ARI and label-matched clustering accuracy are reported; the overall best value is shown in bold and the benchmark best value is underlined. Benchmark pairs are ordered by the number of scenarios in which they ranked among the top five by ARI.

| Method | Balanced strong | | | | | | | | Unbalanced strong | | | | | | | |
|---|---|---|---|---|---|---|---|---|---|---|---|---|---|---|---|---|
| | $n = 500$ | | $n = 1000$ | | $n = 2000$ | | $n = 4000$ | | $n = 500$ | | $n = 1000$ | | $n = 2000$ | | $n = 4000$ | |
| | ARI | Acc. | ARI | Acc. | ARI | Acc. | ARI | Acc. | ARI | Acc. | ARI | Acc. | ARI | Acc. | ARI | Acc. |
| **SMORE** | 0.758 | 0.8072 | **0.847** | **0.9007** | **0.807** | **0.8404** | **0.830** | **0.8711** | **0.917** | **0.9428** | **0.905** | 0.9495 | **0.906** | **0.9500** | **0.911** | **0.9535** |
| PCA + Louvain | **0.781** | 0.800 | 0.782 | 0.800 | 0.770 | 0.803 | 0.762 | 0.797 | 0.859 | 0.906 | 0.707 | 0.770 | 0.689 | 0.752 | 0.676 | 0.734 |
| PCA + GMM | **0.781** | 0.800 | 0.803 | 0.830 | 0.782 | 0.800 | 0.782 | 0.800 | 0.722 | 0.837 | 0.543 | 0.730 | **0.906** | **0.950** | 0.906 | 0.950 |
| PCA + K-means | 0.671 | 0.717 | 0.743 | 0.774 | 0.675 | 0.713 | 0.747 | 0.770 | 0.776 | 0.800 | 0.704 | 0.769 | 0.773 | 0.835 | 0.634 | 0.785 |
| t-SNE + Hierarchical | 0.748 | 0.803 | 0.782 | 0.800 | 0.782 | 0.800 | 0.782 | 0.800 | 0.461 | 0.665 | 0.870 | 0.924 | 0.882 | 0.928 | 0.901 | 0.947 |
| PCA + DBSCAN | 0.779 | **0.816** | 0.782 | 0.801 | 0.782 | 0.800 | 0.782 | 0.800 | 0.890 | 0.940 | **0.905** | **0.950** | **0.906** | **0.950** | 0.906 | 0.950 |
| t-SNE + K-means | 0.697 | 0.763 | 0.753 | 0.772 | 0.693 | 0.720 | 0.659 | 0.702 | 0.378 | 0.551 | 0.684 | 0.742 | 0.848 | 0.891 | 0.765 | 0.805 |
| t-SNE + GMM | 0.711 | 0.760 | 0.782 | 0.800 | 0.548 | 0.620 | 0.548 | 0.620 | 0.429 | 0.656 | 0.826 | 0.894 | 0.362 | 0.620 | 0.795 | 0.882 |
| UMAP + GMM | 0.477 | 0.600 | 0.478 | 0.600 | 0.480 | 0.600 | 0.480 | 0.600 | 0.307 | 0.578 | 0.252 | 0.547 | 0.253 | 0.547 | 0.422 | 0.646 |
| NMF + Louvain | 0.718 | 0.783 | 0.747 | 0.803 | 0.743 | 0.792 | 0.735 | 0.775 | 0.408 | 0.519 | 0.541 | 0.639 | 0.594 | 0.681 | 0.570 | 0.651 |
| UMAP + Hierarchical | 0.477 | 0.600 | 0.478 | 0.600 | 0.480 | 0.600 | 0.480 | 0.600 | 0.438 | 0.654 | 0.420 | 0.645 | 0.420 | 0.645 | 0.422 | 0.646 |

| Method | Balanced weak | | | | | | | | Unbalanced weak | | | | | | | |
|---|---|---|---|---|---|---|---|---|---|---|---|---|---|---|---|---|
| | $n = 500$ | | $n = 1000$ | | $n = 2000$ | | $n = 4000$ | | $n = 500$ | | $n = 1000$ | | $n = 2000$ | | $n = 4000$ | |
| | ARI | Acc. | ARI | Acc. | ARI | Acc. | ARI | Acc. | ARI | Acc. | ARI | Acc. | ARI | Acc. | ARI | Acc. |
| **SMORE** | 0.381 | 0.5510 | 0.414 | 0.5744 | 0.433 | 0.5937 | 0.531 | 0.6377 | **0.553** | **0.7406** | 0.685 | 0.8282 | 0.764 | 0.8730 | 0.790 | 0.8876 |
| PCA + Louvain | **0.642** | **0.730** | **0.719** | **0.778** | 0.742 | **0.802** | 0.752 | **0.802** | 0.516 | 0.646 | 0.670 | 0.750 | 0.679 | 0.745 | 0.673 | 0.734 |
| PCA + GMM | 0.522 | 0.620 | 0.481 | 0.600 | 0.512 | 0.620 | 0.512 | 0.620 | 0.206 | 0.529 | 0.410 | 0.650 | 0.709 | 0.830 | 0.858 | 0.920 |
| PCA + K-means | 0.481 | 0.600 | 0.450 | 0.571 | **0.745** | 0.771 | **0.781** | 0.800 | 0.421 | 0.638 | **0.799** | **0.886** | **0.906** | **0.950** | **0.889** | **0.934** |
| t-SNE + Hierarchical | 0.187 | 0.378 | 0.280 | 0.440 | 0.515 | 0.663 | 0.559 | 0.688 | 0.055 | 0.421 | 0.000 | 0.400 | 0.074 | 0.451 | 0.000 | 0.400 |
| PCA + DBSCAN | 0.209 | 0.399 | 0.210 | 0.400 | 0.210 | 0.400 | 0.210 | 0.400 | 0.156 | 0.481 | 0.161 | 0.500 | 0.161 | 0.500 | 0.161 | 0.500 |
| t-SNE + K-means | 0.217 | 0.400 | 0.402 | 0.558 | 0.514 | 0.658 | 0.634 | 0.745 | 0.094 | 0.438 | 0.136 | 0.450 | 0.000 | 0.400 | 0.000 | 0.400 |
| t-SNE + GMM | 0.082 | 0.280 | 0.238 | 0.400 | 0.487 | 0.610 | 0.544 | 0.668 | 0.000 | 0.400 | 0.000 | 0.400 | 0.189 | 0.514 | 0.103 | 0.461 |
| UMAP + GMM | 0.211 | 0.406 | 0.211 | 0.404 | 0.214 | 0.398 | 0.223 | 0.401 | 0.211 | 0.526 | 0.194 | 0.485 | 0.174 | 0.433 | 0.212 | 0.497 |
| NMF + Louvain | 0.022 | 0.202 | 0.017 | 0.159 | 0.029 | 0.152 | 0.139 | 0.234 | 0.002 | 0.185 | 0.003 | 0.142 | 0.002 | 0.130 | 0.001 | 0.116 |
| UMAP + Hierarchical | 0.242 | 0.385 | 0.216 | 0.395 | 0.205 | 0.400 | 0.209 | 0.400 | 0.144 | 0.349 | 0.135 | 0.340 | 0.170 | 0.411 | 0.233 | 0.545 |

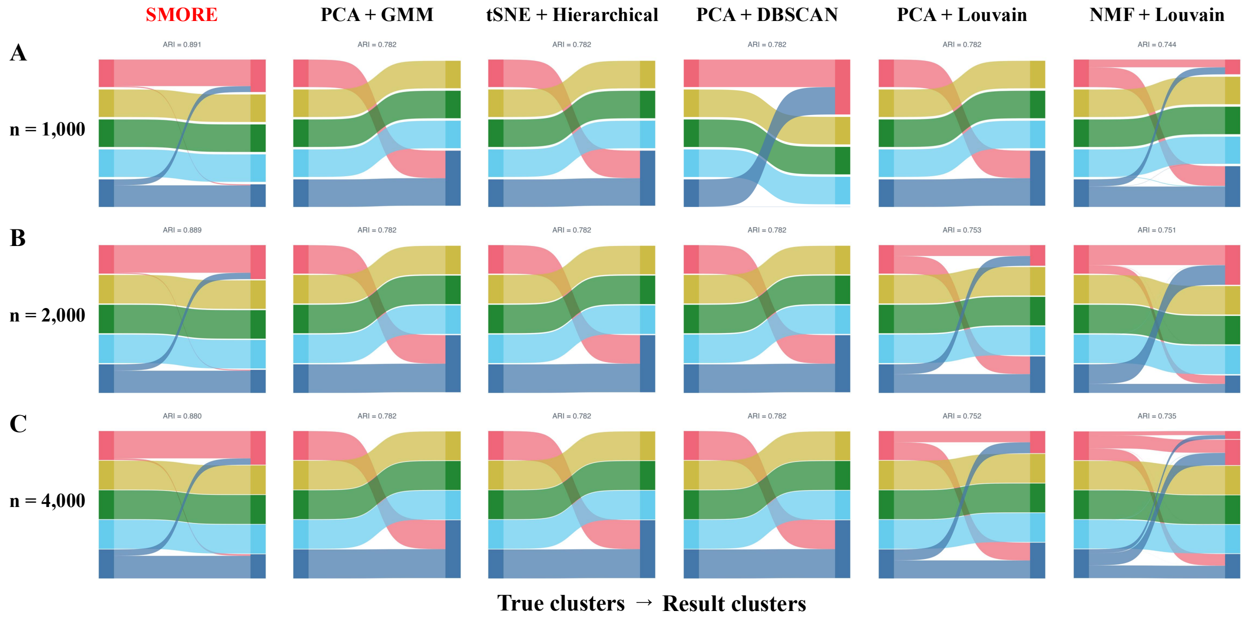

Figure 3: Comparison of cluster-assignment accuracy under the balanced strong-signal simulation setting. **(A)** Sankey diagrams for $n = 1{,}000$ cells. **(B)** Sankey diagrams for $n = 2{,}000$ cells. **(C)** Sankey diagrams for $n = 4{,}000$ cells. Within each panel, diagrams compare the correspondence between the true clusters and those inferred by *SMORE* and five benchmark methods. Colors represent the true clusters, and the width of each flow is proportional to the number of cells shared by the corresponding true and inferred clusters. ARI values are reported above the respective method diagrams.

*SMORE* tends to be conservative and may merge weakly separated groups rather than force a finer partition. However, the method maintained relatively high completeness in these settings, suggesting that it tended to preserve broad latent-group structure even when the separation was insufficient for perfect cell-level assignment. In the unbalanced weak scenarios, performance improved with sample size: mean ARI increased from 0.553 at $n = 500$ to 0.685 at $n = 1000$ and 0.765 at $n = 2000$, with corresponding NMI increasing from 0.563 to 0.757. This pattern indicates that additional cells help stabilize inference when both class imbalance and weak signal are present.

The weaker performance of *SMORE* in the weak-signal settings does not contradict its stronger performance in the model-based simulation reported above, because the two simulations target different sources of clustering difficulty and their absolute ARI values are not directly comparable. In the present simulation, weak signal reduces marginal separation among clusters, while the remaining differences are largely aligned with dominant principal-component directions. This structure favors PCA-based pipelines followed by GMM, DBSCAN, K-means, or Louvain clustering, which can impose relatively sharp partitions in the reduced space. *SMORE*, by contrast, is more conservative when the evidence for distinct latent clusters is weak and may merge poorly separated groups, resulting in lower ARI but relatively preserved completeness. The model-based simulation more closely follows the latent ordinal factor mixture structure assumed by *SMORE*, allowing it to exploit the underlying generative structure directly. The relative strength of PCA-based benchmarks in the present simulation should therefore be interpreted as specific to this stress-test design rather than as a general advantage across data-generating settings.

#### S7.2 Computational runtime

Table 2: Computational runtime in the simulation studies. Runtime is reported in minutes as median [interquartile range]. The conventional-workflow summary pools all recorded dimension-reduction and clustering workflows across the same scenarios and replicates.

| | | Computational runtime | |
|---|---|---|---|
| Simulation design | Cells ($n$) | ORBIT | Benchmark methods |
| Model-based | 500 | 18.1 [17.3–18.4] | 0.26 [0.03–3.23] |
| Model-based | 1,000 | 35.3 [34.6–35.7] | 0.15 [0.11–4.64] |
| Model-based | 2,000 | 71.0 [69.2–72.1] | 0.61 [0.47–3.67] |
| Model-based | 4,000 | 177.3 [176.4–178.5] | 2.59 [2.13–4.37] |
| Distribution-based | 500 | 21.5 [21.2–21.7] | 0.31 [0.03–3.54] |
| Distribution-based | 1,000 | 42.9 [42.1–43.7] | 0.14 [0.11–3.54] |
| Distribution-based | 2,000 | 86.8 [85.2–88.7] | 0.57 [0.50–3.67] |
| Distribution-based | 4,000 | 175.9 [172.1–180.6] | 2.45 [2.13–4.10] |

The computational runtime of *SMORE* and the conventional dimension-reduction and clustering workflows under the two simulation designs is summarized in Supplementary Table 2. As expected, fitting the full Bayesian latent-variable clustering model by MCMC required substantially more computation than the conventional two-stage workflows. Nevertheless, *SMORE* was computationally feasible for datasets containing up to 4,000 cells and 5,000 features.

### S8 Real data analysis

#### S8.1 Data preparation detail

Real methylation data were preprocessed to ensure data quality and compatibility with both the proposed ordinal modeling framework and the benchmark clustering pipelines. The preprocessing choices were motivated by the characteristics of single-cell methylation data, including sparse coverage, boundary-inflated methylation proportions, and the naturally ordered interpretation of methylation states, rather than by method-specific optimization. The same processed ordinal matrix was used for all methods to avoid giving *SMORE* an advantage through a separate preprocessing pipeline.

The primary real-data analysis focused on the lung dataset, additional analyses of the are PBMC and primary motor cortex datasets. For each dataset, the available methylated-count matrix and coverage matrix were used to construct methylation profiles over fixed genomic bins, with rows corresponding to genomic regions and columns corresponding to cells.

**Origin of reference annotations** The cell labels used for external evaluation were obtained from the metadata deposited with the Human Body Single-Cell Atlas of 3D Genome Organization and DNA Methylation (Zhou et al., 2026). In the source study, label construction combined algorithmic clustering with manual assessment and reference-guided biological annotation. Cells were first clustered using joint DNA-methylation and chromatin-contact embeddings, with the embedding dimensions used for within-major-type clustering manually examined to ensure that the resulting partitions were stable. Initial consensus Leiden clusters were then iteratively merged according to their separability across the two molecular modalities. The resulting clusters were assigned biological cell-type annotations by considering tissue of origin and their correspondence with annotated scRNA-seq cell types, supported by differential gene expression, differentially methylated regions, and differential chromatin loops. Thus, the deposited labels were not generated by a fully automated clustering procedure alone, but incorporated manual assessment and multiple sources of biological evidence. Nevertheless, because the labels were assigned at the cluster level rather than independently measured for each cell, we treat them as curated source-study reference annotations rather than definitive ground-truth labels.

**Methylation level computation** For each genomic bin and cell, the methylation level was computed as the ratio of methylated counts to total coverage. Entries for which both the methylated count and coverage were zero were treated as missing values, because such entries do not contain informative methylation measurements. This produced a continuous methylation proportion matrix with entries in $[0, 1]$ before downstream discretization.

**Quality control** To reduce noise caused by excessive sparsity, filtering was applied at both the cell and feature levels. Cells with missing rate greater than $75\%$ were excluded, and genomic bins with missing rate greater than $70\%$ were removed. This filtering step retained cells and genomic regions with sufficient observed methylation information for reliable clustering analysis.

**Imputation via kernel density estimation** Missing values were imputed using a kernel density estimation (KDE)-based approach applied column-wise (Parzen, 1962), separately for each cell. This choice was intended to preserve cell-specific methylation distributions rather than borrowing a single global distribution across all cells. Such a strategy is appropriate for single-cell methylation data because cells can differ substantially in their overall methylation profiles and coverage patterns.

For each cell, when sufficient observed values were available, a kernel density estimate was fitted to the observed methylation proportions on $[0, 1]$, and missing entries were imputed by sampling from the estimated density. If too few observed values were available for reliable density estimation, a simple fallback rule based on the global median was used to avoid unstable imputation. Imputed values were truncated to $[0, 1]$. This imputation step was used only to obtain a complete methylation matrix for downstream comparison; it was applied before the common ordinal transformation and was not tuned separately for the proposed method.

**Ordinal transformation** Although continuous imputed methylation matrices were generated during preprocessing, all real-data analyses used an ordinal representation as the common input. The imputed methylation proportions were discretized using the same five-level transformation defined for the distribution-mixture simulation in Section S7. This transformation retains more information than a binary methylated/unmethylated representation while reducing sensitivity to small continuous differences that may arise from sparse or heterogeneous sequencing coverage.

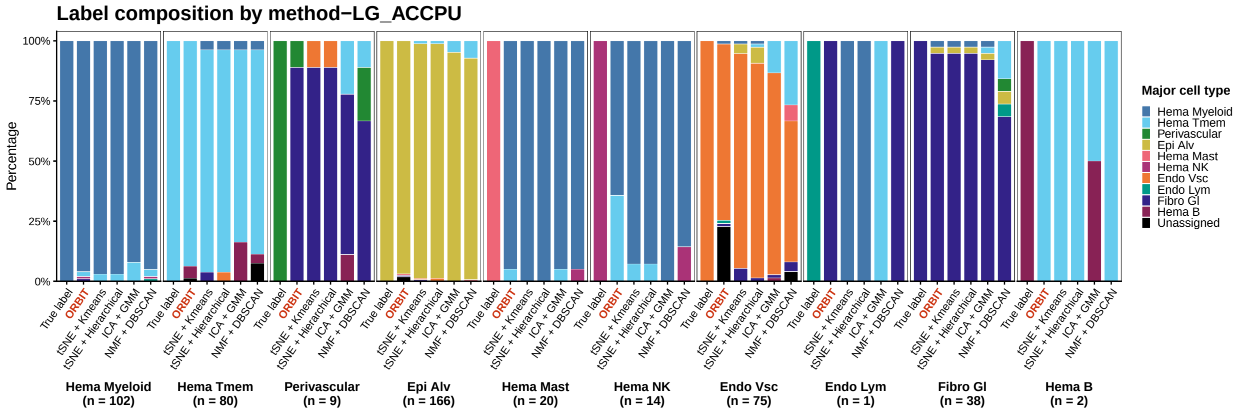

Figure 4: Percentage composition of reference cell types across clustering results for the lung dataset. Bars are grouped by reference cell type and compare the proposed method with the selected benchmark methods. Colors indicate cell types, and gray segments represent unassigned cells. Sample sizes for the reference cell-type groups are shown below the corresponding bars.

The resulting ordinal matrix was used as input for both *SMORE* and all benchmark pipelines, ensuring that all methods were evaluated using the same processed data.

**Variance filtering** Features with zero variance were removed prior to downstream analysis, as they do not contribute to clustering or dimensionality reduction and may lead to numerical instability (e.g., in principal component analysis).

#### S8.2 Additional result-lung

The percentage-normalized composition provides a complementary view to the frequency-based results presented in the main text. By removing differences in absolute cell-type abundance, this representation highlights the purity of the inferred clusters associated with each reference major cell type. Across methods, several major cell types, including Hema Myeloid, Hema Tmem, Epi Alv, Hema Mast, and Endo Lym, were represented by clusters dominated by the corresponding reference labels, indicating that these populations were comparatively well separated. In contrast, more heterogeneous compositions were observed for Perivascular, Endo Vsc, Fibro Gl, and some of the smaller cell populations, suggesting partial overlap or reassignment among closely related methylation profiles. The percentage compositions of very rare cell types, particularly Hema B ($n = 2$), should be interpreted cautiously because the assignment of a single cell can produce a

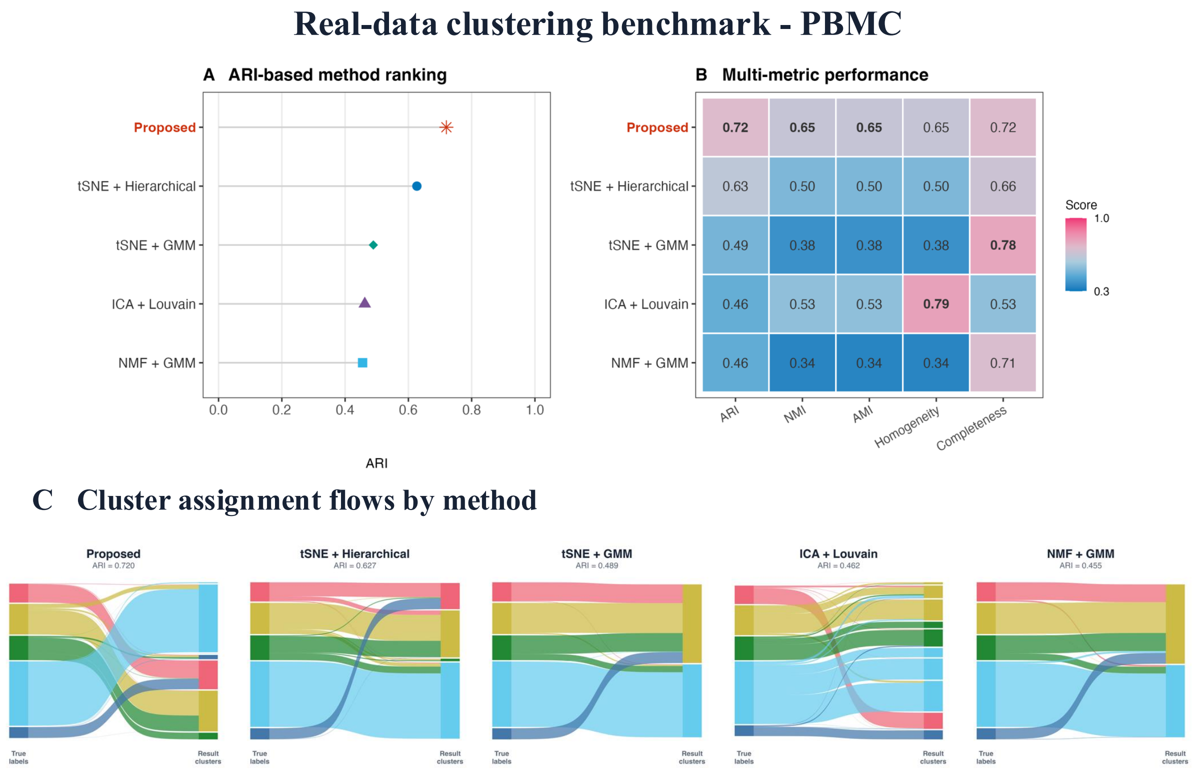

Figure 5: Real-data clustering benchmark on the PBMC dataset ($n = 1{,}975$). **(A)** ARI-based ranking of the proposed method and the four top-performing benchmark methods. **(B)** Comparison across ARI, NMI, AMI, homogeneity, and completeness, with the highest value for each metric shown in bold. **(C)** Sankey diagrams showing cell-assignment flows from the reference cell-type labels to the inferred clusters for each method. Colors represent reference cell types and are propagated through the corresponding assignment flows.

large proportional change. Together with the frequency-based visualization, these results show that differences between methods arise not only from the recovery of abundant populations but also from their ability to maintain cluster purity for less distinct or low-frequency cell types.

#### S8.3 Result-PBMC dataset

We next evaluated the proposed method on the processed PBMC dataset. After preprocessing, the dataset contained 1,975 cells from five annotated reference groups: c0, c1, c15, c27 and c7, comprising 145, 852, 406, 250 and 322 cells, respectively. The marked differences in group size provided an additional test of clustering performance under class imbalance. All methods were applied to the same processed ordinal methylation matrix, and adjusted Rand index (ARI) was

Table 3: Clustering performance of the top-10 benchmark method pairs and the proposed method on the PBMC dataset ($n = 1,975$ cells). **Bold**: proposed method. Underline: best-performing benchmark per metric.

| Method | ARI | NMI | AMI | Hom. | Comp. |
|---|---|---|---|---|---|
| **SMORE** | **0.720** | **0.649** | **0.648** | **0.649** | **0.717** |
| t-SNE + Hierarchical | 0.627 | 0.500 | 0.499 | 0.500 | 0.665 |
| t-SNE + GMM | 0.489 | 0.376 | 0.376 | 0.376 | 0.781 |
| ICA + Louvain | 0.462 | 0.532 | 0.530 | 0.793 | 0.532 |
| NMF + GMM | 0.455 | 0.340 | 0.340 | 0.340 | 0.707 |
| ICA + K-means | 0.454 | 0.361 | 0.361 | 0.361 | 0.754 |
| t-SNE + K-means | 0.442 | 0.529 | 0.528 | 0.529 | 0.551 |
| ICA + GMM | 0.441 | 0.329 | 0.328 | 0.329 | 0.683 |
| PCA + GMM | 0.428 | 0.320 | 0.320 | 0.320 | 0.665 |
| PCA + Louvain | 0.424 | 0.533 | 0.531 | 0.865 | 0.533 |
| NMF + K-means | 0.405 | 0.345 | 0.344 | 0.345 | 0.727 |

used as the primary evaluation metric.

*SMORE* achieved the highest ARI of 0.720, compared with 0.627 for the strongest benchmark, t-SNE followed by hierarchical clustering (Figure 5A,B and Table 3). It also achieved the highest NMI and AMI among the evaluated methods. Although PCA followed by Louvain clustering attained the highest homogeneity and t-SNE followed by GMM attained the highest completeness, both showed substantially lower overall agreement with the reference labels. These results indicate that *SMORE* provided the strongest and most balanced agreement across the primary and information-based clustering metrics.

The Sankey diagrams further illustrate these differences (Figure 5C). The proposed method largely preserved the dominant c1 group while also separating substantial fractions of the c15, c27 and c7 groups. By contrast, several benchmark pipelines produced coarser partitions in which multiple reference groups converged into the same inferred cluster. The proposed method inferred seven clusters for the five reference groups, indicating some subdivision relative to the available annotations. Nevertheless, its higher ARI suggests that the resulting partition remained more concordant with the reference structure than the benchmark solutions.

The composition analysis showed that most cells in the largest reference group, c1, were assigned consistently by the proposed method (Figure 6). The c15, c27 and c7 groups also retained

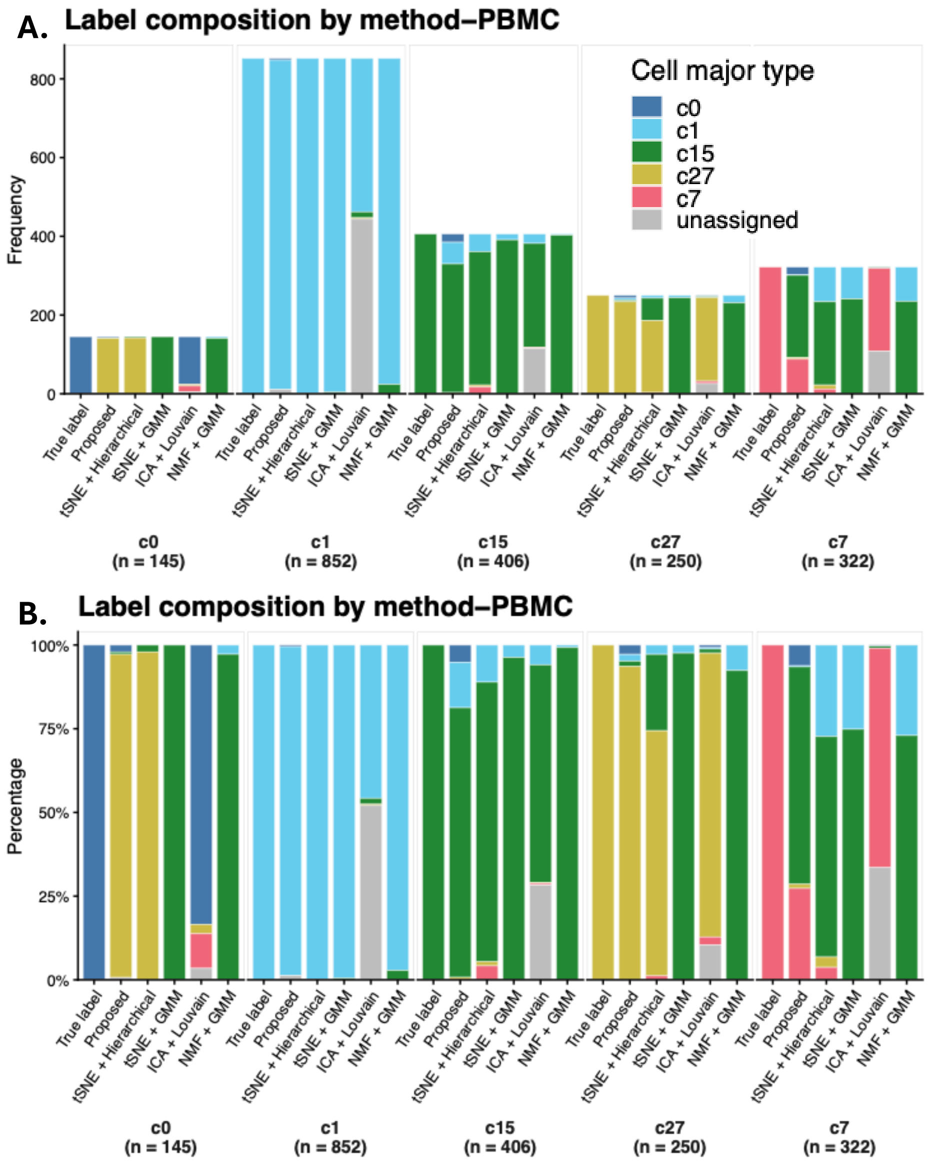

Figure 6: Composition of reference major cell types across clustering results for the PBMC dataset. **(A)** Cell frequencies and **(B)** percentage-normalized compositions for each reference major cell type across SMORE and the selected benchmark methods. Colors indicate the reference major cell types, gray segments represent unassigned cells, and the corresponding reference-group sample sizes are shown below the panels.

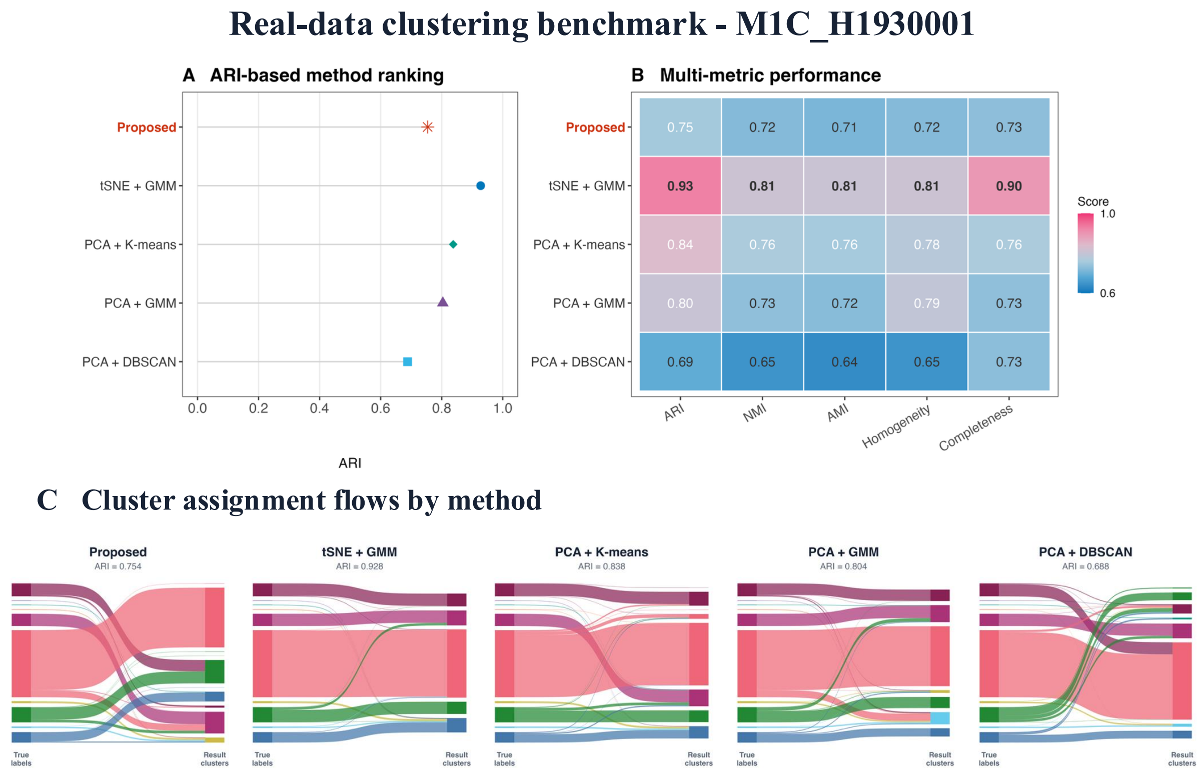

Figure 7: Real-data clustering benchmark on the primary motor cortex dataset ($n = 540$). **(A)** ARI-based ranking of the proposed method and the four top-performing benchmark methods. **(B)** Comparison across ARI, NMI, AMI, homogeneity, and completeness, with the highest value for each metric shown in bold. **(C)** Sankey diagrams showing cell-assignment flows from the reference cell-type labels to the inferred clusters. Colors represent reference cell types and their corresponding assignment flows.

clear dominant matched components, although some mixing remained. Errors were more pronounced for the smallest c0 group, which was not cleanly separated by the proposed method. This pattern indicates that the remaining disagreement was concentrated primarily in the smaller reference groups rather than being distributed uniformly across the dataset. As the frequency-scale visualization is strongly influenced by the unequal reference-group sizes, the quantitative comparisons in Table 3, rather than the absolute bar heights, were used to assess overall clustering performance.

Table 4: Clustering performance of the proposed method and the ten highest-ranked benchmark pipelines on the primary motor cortex dataset ($n = 540$ cells). Benchmark pipelines are ordered by adjusted Rand index (ARI). NMI, normalized mutual information; AMI, adjusted mutual information; Hom., homogeneity; Comp., completeness. Boldface identifies the proposed method, and underlining identifies the best-performing benchmark pipeline for each metric.

| Method | ARI | NMI | AMI | Hom. | Comp. |
|---|---|---|---|---|---|
| **SMORE** | 0.754 | 0.721 | 0.711 | 0.721 | 0.730 |
| t-SNE + GMM | **0.928** | **0.810** | **0.805** | <u>0.810</u> | **0.896** |
| PCA + K-means | <u>0.838</u> | <u>0.763</u> | <u>0.756</u> | 0.779 | <u>0.763</u> |
| PCA + GMM | 0.804 | 0.732 | 0.724 | 0.793 | 0.732 |
| PCA + DBSCAN | 0.688 | 0.647 | 0.635 | 0.647 | 0.729 |
| NMF + K-means | 0.686 | 0.628 | 0.618 | 0.679 | 0.628 |
| NMF + DBSCAN | 0.648 | 0.649 | 0.641 | 0.649 | 0.654 |
| ICA + DBSCAN | 0.624 | 0.612 | 0.593 | 0.612 | 0.625 |
| ICA + GMM | 0.590 | 0.657 | 0.647 | 0.797 | 0.657 |
| NMF + GMM | 0.550 | 0.607 | 0.595 | 0.767 | 0.607 |
| t-SNE + Hierarchical | 0.523 | 0.642 | 0.631 | **0.833** | 0.642 |

#### S8.4 Result-primary motor cortex

We further evaluated the proposed method on the primary motor cortex single-cell methylation dataset. After preprocessing, the dataset contained 540 cells assigned to ten reference groups. The reference-group sizes were highly unequal: c14 was the dominant group with 296 cells, whereas c19, c27 and c29 contained only two, two and one cell, respectively. This dataset therefore provided a challenging setting for assessing clustering performance in the presence of pronounced class imbalance and several rare cell populations.

*SMORE* achieved an ARI of 0.754 but was not the highest-performing method on the primary motor cortex dataset (Figure 7A,B and Table 4). t-SNE followed by GMM achieved the highest ARI of 0.928 and also led the comparison in NMI, AMI, and completeness. Two PCA-based pipelines also attained higher ARI than *SMORE*, indicating that the strongest embedding-based benchmarks aligned particularly well with the reference structure in this dataset.

The MDS and label-composition analyses help explain the performance differences between the proposed method and t-SNE combined with GMM (Figures 8 and 9). Although the reference annotations contained ten groups, the five larger groups comprised 520 of the 540 cells (96.3%). t-SNE combined with GMM inferred five clusters that closely recovered these dominant populations.

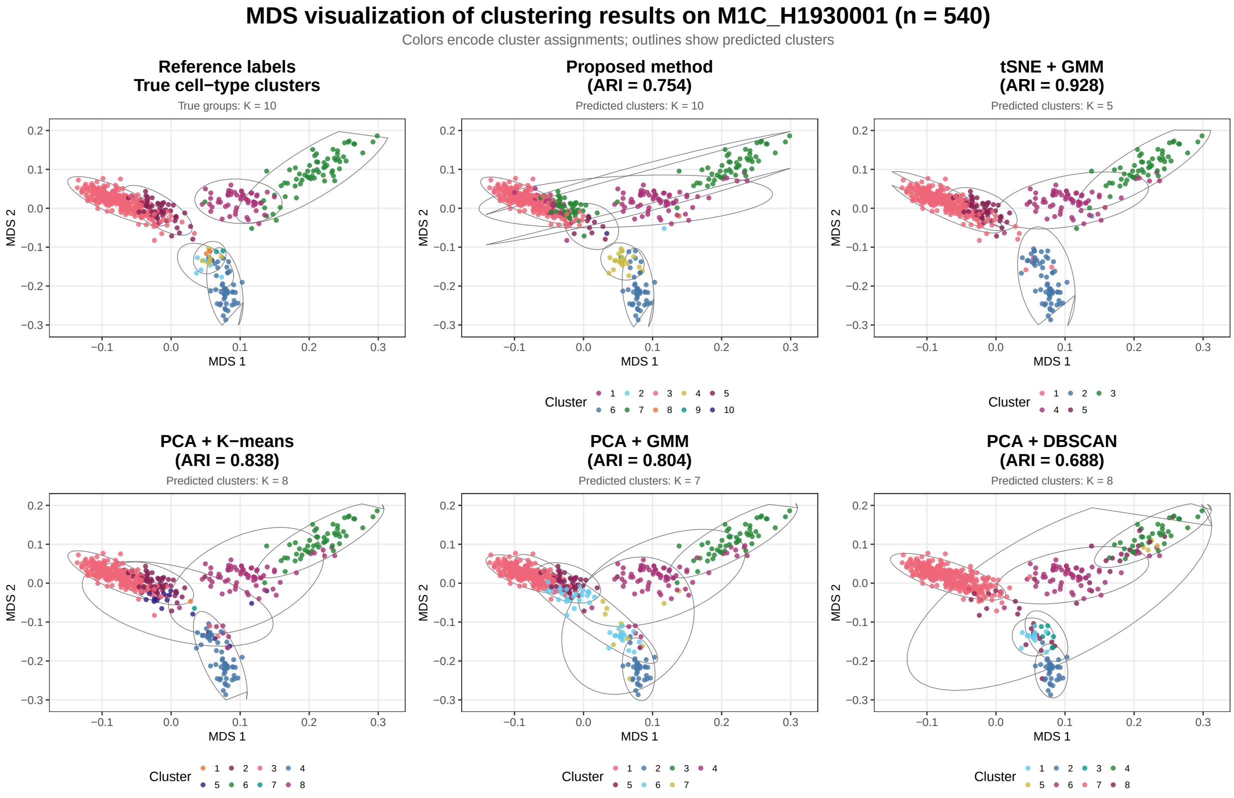

Figure 8: MDS visualization of clustering results on the primary motor cortex dataset. The first panel shows the reference cell-type labels, followed by the proposed method and the four top-performing benchmark methods. Point colors indicate cluster assignments, while ellipses outline the inferred clusters. ARI values and the numbers of predicted clusters are reported above the corresponding method panels.

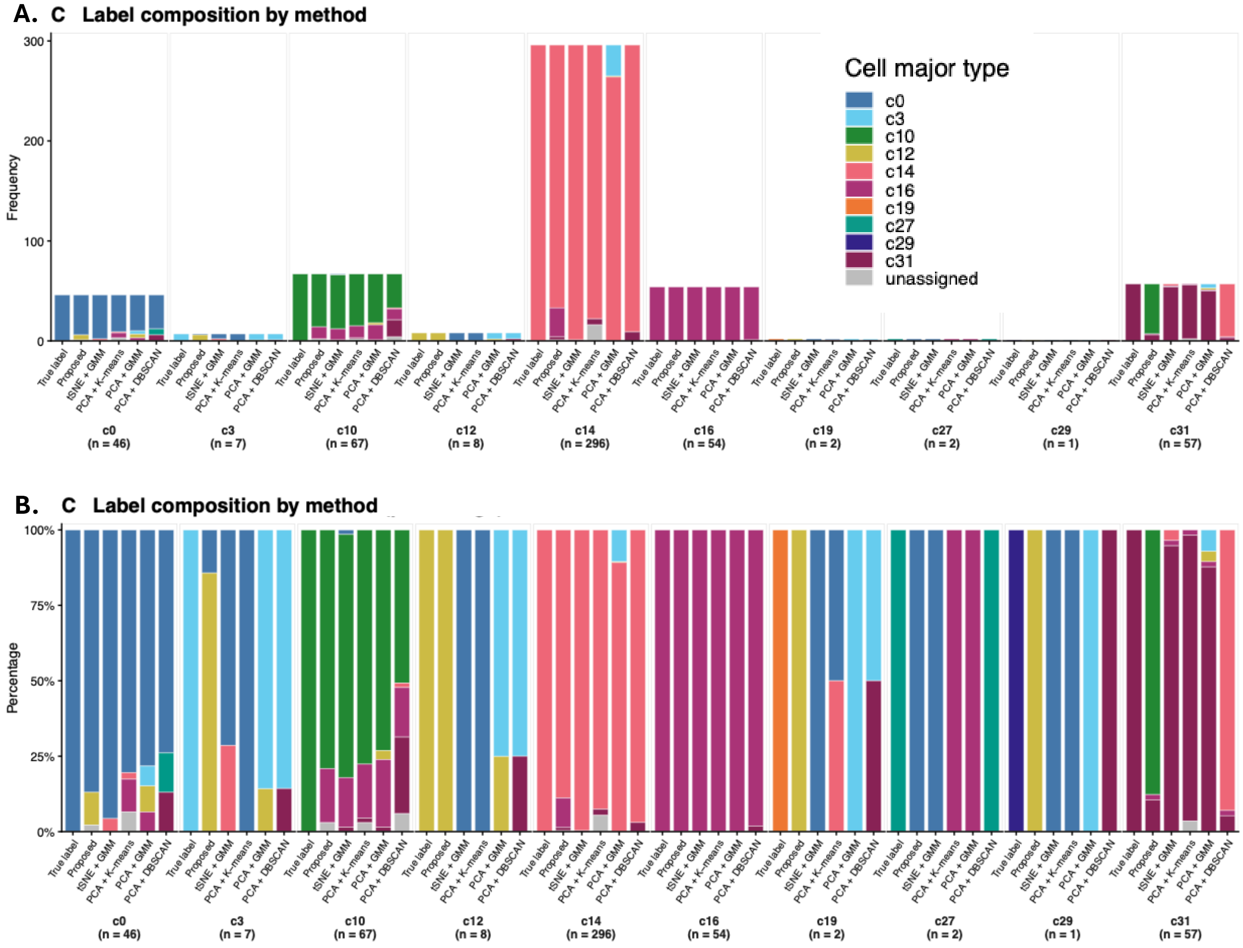

Figure 9: Composition of reference major cell types across clustering results for the primary motor cortex dataset. **(A)** Cell frequencies and **(B)** percentage-normalized compositions for each reference major cell type across SMORE and the selected benchmark methods. Colors indicate the reference major cell types, gray segments represent unassigned cells, and the corresponding reference-group sample sizes are shown below the panels.

Because ARI is based on pairwise agreement, accurate recovery of the larger groups contributed substantially to its high ARI, whereas merging several rare groups had a comparatively limited effect on the overall score. The clear separation of the larger populations in the MDS representation is consistent with the strong performance of this embedding-based pipeline on the primary motor cortex dataset.

The proposed method inferred ten clusters without requiring the cluster number to be specified in advance, matching the number of reference groups. It recovered dominant matched components for several larger groups, including c0, c10, c14 and c16, while retaining a finer-grained partition than t-SNE combined with GMM. However, some of these subdivisions did not align cleanly with the reference annotations, particularly for c31 and the smallest groups, resulting in lower overall agreement with the reference labels. Thus, the proposed method preserved more components but achieved less accurate overall correspondence with the available reference labels, whereas t-SNE combined with GMM favored a coarser partition dominated by the five abundant populations.

The M1C results therefore identify a setting in which the proposed method did not outperform the strongest benchmark, while showing that differences in class abundance and cluster granularity can substantially affect method rankings. Together with the lung and PBMC analyses, these findings indicate that clustering performance depends on both the distributional structure of the data and the relative abundance of the underlying cell populations.

#### S8.5 Sensitivity analysis for latent dimension

In supplementary Table 5, we evaluated the sensitivity of MFM to the tuning parameter $q$ on three real datasets by setting $q \in \{5, 10, 15, 20\}$ while keeping the remaining unchanged. The optimal value of $q$ was dataset dependent. For PBMC and primary motor cortex datasets, the highest ARI was obtained at $q = 5$. However, their ARI values varied over relatively narrow ranges of 0.019 and 0.051, respectively, suggesting that the clustering results for these two datasets were relatively more robust to the choice of $q$. In contrast, LG-ACCPU exhibited substantially greater sensitivity: its ARI increased to 0.830 at $q = 10$, compared with 0.610, 0.544, and 0.632 at $q = 5$, 15, and 20, respectively. Similar patterns were observed for NMI, AMI, FMI, and clustering accuracy. Consequently, although $q = 10$ produced the highest average performance across the three datasets, this result was driven primarily by its pronounced improvement on LG-ACCPU rather than by uniformly superior performance across all datasets.

These results suggest that increasing $q$ does not lead to a monotonic improvement in clustering performance. A larger $q$ provides a more flexible latent representation, which may be beneficial when the underlying cellular structure is relatively complex, as appears to be the case for LG-ACCPU. However, once sufficient flexibility has been reached, further increasing $q$ may introduce redundant dimensions or additional estimation variability without improving separation between cell populations. This may explain why the optimal option of $q$ was dataset dependent.

Within each dataset, runtime generally increased as $q$ increased. From $q = 5$ to $q = 20$, runtime increased by 14.8 percent, 13.7 percent, and 15.2 percent for LG-ACCPU, PBMC, and primary motor cortex datasets, respectively, with a broadly monotonic pattern. This increase is expected because a larger $q$ expands the latent parameter space explored during model fitting. In contrast, clustering accuracy did not improve monotonically with $q$, indicating diminishing returns from unnecessarily large values of $q$. A moderate value may therefore be used as a practical default, with dataset-specific tuning when clustering performance shows greater sensitivity. Absolute runtimes under the default setting and comparisons with benchmark workflows are reported separately in Supplementary Section S8.6.

Table 5: Sensitivity of SMORE to the tuning parameter $q$ on three real datasets. Comp. denotes completeness, Hom. denotes homogeneity, Acc. denotes clustering accuracy. Runtime is reported in hours. Boldface identifies the best similarity or accuracy value within each dataset; runtime is not used for boldface selection.

| Dataset | $n$ | $q$ | ARI | NMI | AMI | FMI | Comp. | Hom. | Acc. | Runtime (h) |
|---|---|---|---|---|---|---|---|---|---|---|
| Lung | 507 | 5 | 0.610 | 0.592 | 0.586 | 0.723 | **0.843** | 0.592 | 0.690 | 2.06 |
| | | 10 | **0.830** | **0.770** | **0.763** | **0.866** | 0.836 | **0.770** | **0.860** | 2.16 |
| | | 15 | 0.544 | 0.519 | 0.505 | 0.685 | 0.781 | 0.519 | 0.663 | 2.26 |
| | | 20 | 0.632 | 0.642 | 0.627 | 0.735 | 0.799 | 0.642 | 0.706 | 2.37 |
| PBMC | 1,975 | 5 | **0.733** | **0.664** | **0.663** | **0.814** | 0.739 | **0.664** | 0.758 | 8.16 |
| | | 10 | 0.720 | 0.649 | 0.648 | 0.806 | 0.717 | 0.649 | 0.754 | 8.80 |
| | | 15 | 0.715 | 0.596 | 0.594 | 0.806 | 0.741 | 0.596 | 0.732 | 8.87 |
| | | 20 | 0.719 | 0.661 | 0.659 | 0.805 | **0.752** | 0.661 | **0.764** | 9.28 |
| Primary motor cortex | 540 | 5 | **0.772** | 0.739 | 0.732 | **0.848** | **0.785** | 0.739 | **0.796** | 2.20 |
| | | 10 | 0.754 | 0.721 | 0.711 | 0.835 | 0.730 | 0.721 | 0.785 | 2.71 |
| | | 15 | 0.735 | **0.750** | **0.740** | 0.822 | 0.754 | **0.750** | 0.783 | 2.42 |
| | | 20 | 0.721 | 0.676 | 0.663 | 0.812 | 0.676 | 0.712 | 0.767 | 2.54 |

#### S8.6 Computational runtime

Table 6: Computational runtime on the three real single-cell DNA methylation datasets. Proposed-method runtime is the elapsed time for the recorded Bayesian model fit. For context, the conventional-workflow column reports the median [minimum–maximum] elapsed time across all recorded dimension-reduction and clustering workflows for each dataset. Times are in minutes.

| | | | Computational runtime | |
|---|---|---|---|---|
| Dataset | Cells ($n$) | Genomic bin count | ORBIT | Benchmark methods |
| Lung | 507 | 15,209 | 129.69 | 0.34 [0.04–90.10] |
| PBMC | 1,975 | 15,239 | 527.71 | 1.25 [0.48–92.25] |
| Primary motor cortex | 540 | 15,430 | 162.52 | 0.37 [0.04–92.36] |

Supplementary Table 6 reports the computational runtime on the three real datasets under the default setting of $q = 10$. The proposed method required longer runtime than conventional dimension-reduction and clustering workflows, reflecting the additional computational cost of joint Bayesian inference for the latent representation, cluster assignments, and number of clusters. As shown in the sensitivity analysis, runtime also increased moderately with $q$ because larger values expand the latent parameter space explored during model fitting. The reported times should therefore be interpreted as computational-cost summaries for the selected model setting rather than as comparisons between equivalent inferential procedures.

### S9 Additional analysis of results

**MCMC diagnostics for simulation studies and real-data analyses** To examine the behavior of the MCMC sampler after burn-in, we inspected retained posterior traces from representative simulation and real-data analyses. Specifically, we monitored the 10 latent-dimensional mean parameters and corresponding diagonal variance parameters for one reference cluster, together with the ARI of the sampled partitions relative to the true simulation labels or reference cell-type annotations. Across the representative analyses, the parameter traces generally remained within stable ranges over the retained iterations, although the degree of fluctuation and gradual movement varied across parameters and datasets. The ARI traces likewise remained within dataset-specific ranges, suggesting that the sampled partitions maintained broadly consistent agreement with the

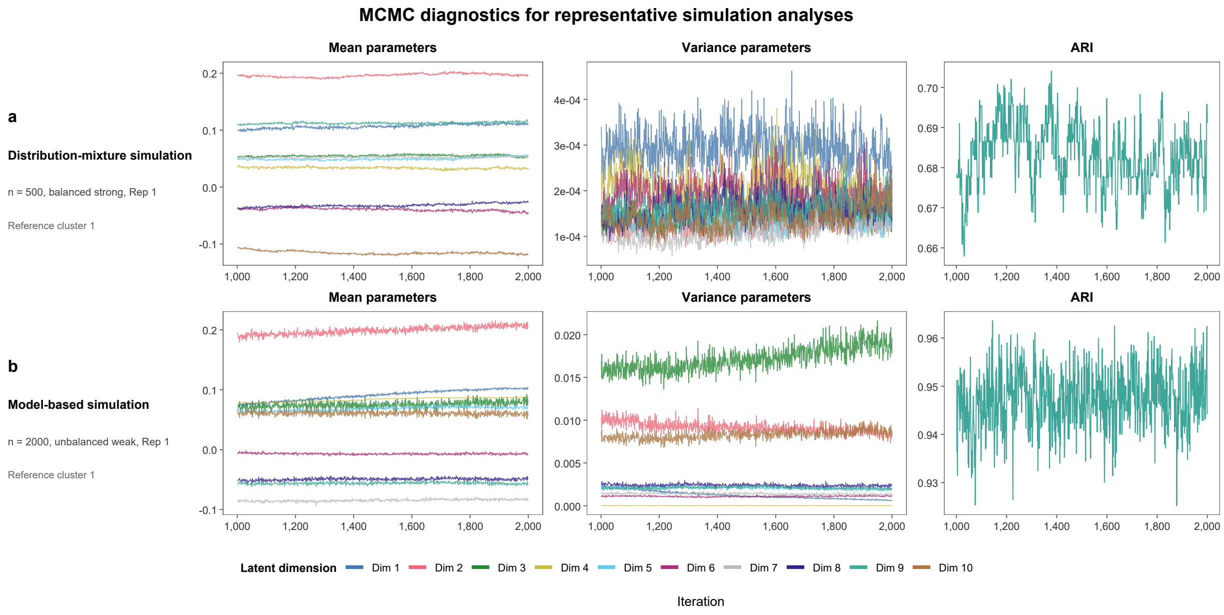

Figure 10: MCMC diagnostics for representative simulation analyses. **(A)** Trace plots for the first replicate of the distribution-mixture simulation with $n = 500$ cells under the balanced strong-signal setting. **(B)** Trace plots for the first replicate of the model-based simulation with $n = 2{,}000$ cells under the unbalanced weak-signal setting. For each analysis, one reference cluster was monitored across the retained posterior samples. The first two columns show the 10 cluster-specific mean parameters and the corresponding diagonal covariance parameters, respectively, and the third column shows the adjusted Rand index relative to the true cluster labels. The parameter traces assess the sampling stability of the cluster-specific model parameters, whereas the ARI trace summarizes partition recovery and stability.

corresponding labels over the retained posterior draws. Figures 10 and 11 present the simulation and real-data results, respectively.

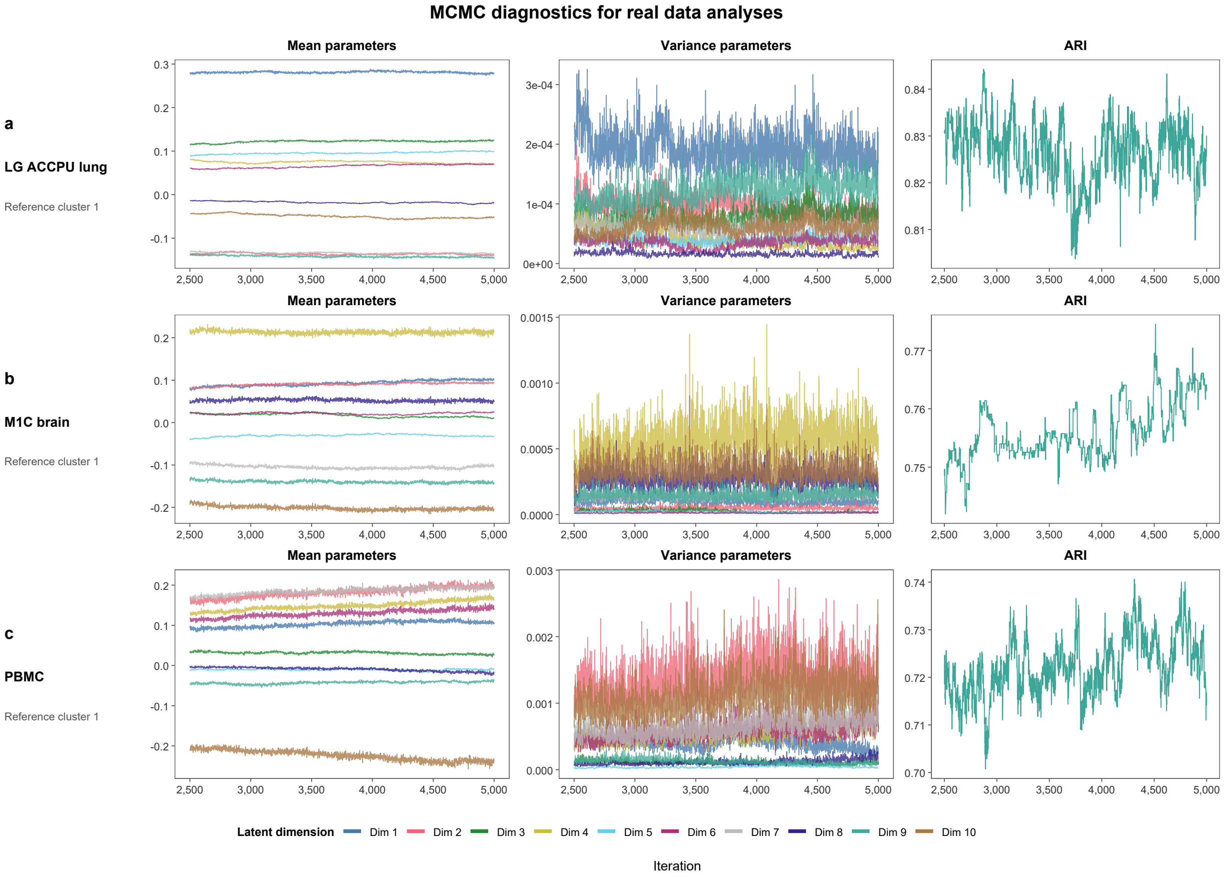

Figure 11: MCMC diagnostics for the real data analyses. Trace plots are shown for the LG ACCPU lung, M1C brain, and PBMC datasets. For each dataset, a single reference cluster was selected, and the first two panels display the retained MCMC traces of its 10 latent dimensional mean parameters and corresponding diagonal variance parameters, respectively. The final panel shows the adjusted Rand index relative to the reference cell type annotations across retained posterior draws. The parameter traces are used to assess stability of the sampled cluster specific model parameters, whereas ARI is presented as a measure of partition stability and agreement with the reference annotations.

Parzen, E. (1962). On estimation of a probability density function and mode. *The Annals of Mathematical Statistics*, *33*(3), 1065–1076.

Zhou, J., Wu, Y., Liu, H., Tian, W., Castanon, R. G., Bartlett, A., … Ecker, J. R. (2026). Human body single-cell atlas of 3D genome organization and DNA methylation. *Science*, *393*(6809), eadx0673. doi: 10.1126/science.adx0673
